# Engineering precision zebrafish alleles of human disease

**DOI:** 10.1101/2025.05.18.654701

**Authors:** Holly R. Thomas, Bradley K. Yoder, Matthew S. Alexander, John M. Parant

**Affiliations:** Department of Cell, Developmental and Integrative Biology, Heersink School of Medicine, University of Alabama at Birmingham, AL, USA; Department of Pediatrics, Division of Neurology at University of Alabama at Birmingham Heersink School of Medicine and Children’s of Alabama, AL, USA

**Keywords:** Precision animal models, Zebrafish, CRIPSR/Cas9, Genome Editing, Oligo directed homology directed repair (HDR), Inference of CRISPR Edits (ICE), or high-resolution melting curve analysis (HRMA)

## Abstract

Animal models of human diseases are an essential component of understanding disease pathogenesis and serve as preclinical models for therapeutic evaluation. Recently human patient genome sequencing has defined unique patient variants that result in disease states with different phenotypes than those observed with null alleles. The UAB Center for Precision Animal Modeling (CPAM) serves to analyze patient variant pathogenicity and disease mechanisms through the generation of animal models. We have optimized a zebrafish gene editing platform to successfully generate 11 patient variants (first round: NF1 R1276Q, NF1 G484R, VMA21 G55V, SPOP D144N, SGO1 K23E, Pex10 H310D, and FKRP C318Y; second round: NF1 R681*, NF1 M992del, P53 R175H, and PKD2 L656W) and 1 research allele (*p53* K120R). We used CRISPR/Cas9 guide directed cleavage along with single-stranded oligodeoxynucleotide (ssODNs) repair templates to generate these models. We evaluated multiple oligo orientations and sizes, but did not find a unified consensus orientation or size that significantly impacted efficiency, emphasizing the need to empirically evaluate multiple variations for the best homology directed repair (HDR) rate. We determined PCR amplicon Next Generation Sequencing (NGS) evaluation of HDR efficiency at the F0 embryo level is best for determining the ideal guide and oligo combination. Further NGS evaluation of DNA from progeny from F0s (germline derived), not F0 biopsy DNA, is essential to identify germline transmitting founders. Surprisingly we find that most founders exhibit a *jackpot* effect in the germ line but not in the somatic tissue. We found NGS superior to using ICE (Inference of CRISPR Edits) for determining HDR frequency. When applicable, allelic-specific PCR or allelic specific restriction digestion can be used to genotype mutation carrying F1 generation animals, however we demonstrated that false positives occur. Further, we successfully used high resolution melting curve analysis (HRMA) to differentiate and identify F1 animals with patient variants.

## INTRODUCTION

Advances in genome sequencing and mapping disease associated variants from whole genome or whole exon sequencing have revolutionized our understanding of disease-causing genes and alleles. Surprisingly many of the disease-causing genes are not genes we would expect to cause these distinct phenotypes. For example, the K23E variants in SGO1 cause a disease called Chronic Atrial and Intestinal Dysrhythmia (CAIDS)^1^. This disease is associated with heart rhythm abnormality and impaired rhythmic muscle contractions that move food through the intestines. SGO1 is considered an essential gene for maintaining sister chromatid cohesion during mitosis, and mouse and zebrafish knockouts of this gene are early embryonic lethal; leaving the question of why does the K23E variant result in a unique disease phenotype ^2^. The UAB Center for Precision Animal Modeling (CPAM) is dedicated to the generation of animal models that carry patient variants. These patient mimetic models are useful for understanding the molecular pathology of the disease but also serve as preclinical models for therapeutic evaluation or screening^3^. Zebrafish is a good vertebrate model of human disease and is therefore one of the organisms that the CPAM has chosen to generate patient mimetic models ^4-10^.

CRISPR/Cas9 genome editing has revolutionized the genome editing capabilities in many organisms including zebrafish. We have established a highly efficient platform for generation of gene knockout alleles using CRISPR/Cas9 in conjunction with high resolution melting curve analysis (HRMA) to identify mutant allele carrying animals ^11^. These null alleles have been very useful in modeling a number of loss-of-function diseases including neurobehavioral, neurodevelopmental, ciliopathies, osteogenesis, cancer, scoliosis, and birth defects ^12-23^. However, patient variant generation or missense alteration has been less well developed in zebrafish. A variety of papers have demonstrated the generation of nucleotide alterations using double stranded DNA ^24-26^. While others have focused on using single-stranded oligodeoxynucleotides (ssODNs) based HDR ^27-38^. The ssODN is simpler since it does not require any cloning steps. Interestingly there are numerous conflicting papers on the best orientation of the ssODN. Some suggest the opposite strand to the PAM, while others emphasize asymmetrical oligos, and multiple papers suggest varying ideal oligo lengths ^28-30,32-34,37,39-42^. However, these studies often draw conclusions based on a limited number of variants generated, thus making it unclear as to what is the best approach to generate zebrafish patient variant carrying animals/ patient mimetic models. We evaluated multiple oligo orientations and sizes, but did not find a unified consensus orientation or size that significantly impacted efficiency, emphasizing the need to empirically evaluate multiple variations at the F0 embryonic stage for the best HDR rate.

Methods to detect the mutated/mutant allele are challenging because often the alterations are just a single nucleotide change, which does not produce a size shift on an agarose gel. Older approaches used single strand conformation polymorphism (SSCP) analysis; however, these are very cumbersome and require radioactivity. The detection method also depends on what level in the production one is studying (F0, F1 or F2). For example, in the Founder (F0) generation, the chimeric animal may carry a patient variant in a sea of CRISPR/Cas9 derived indels. Using traditional sanger sequencing, it is hard to differentiate one specific allele in the combined alleles chromatogram. That said, Inference of CRISPR Edits (ICE) technology essentially evaluates perturbation of the expected chromatograms, along with area under the curve of the perturbation, and predictive algorithms of indels or desired HDR alleles to determine indel and HDR frequencies^43^. High resolution melting curve analysis (HRMA), which detects reduced melting temperature (TM) of heteroduplex molecules in heterozygous samples, can be used to define if there are altered heterozygous sequence but in a chimeric sample cannot differentiate if a specific allele is present ^11^. Allelic specific PCR can be used to define good founders but are limited if allele specific PCR cannot be generated and are often plagued by false positives ^34^. This also has a limit of detection and may not detect changes in low chimeric animals. Allele specific restriction enzyme digestion can be used and incorporates a level of sequence specificity but also has a limited level of detection in low chimeric animals and often require additional silent mutations to generate the restriction site.

The drawback is that these silent mutations could also affect non-coding elements such as splice elements or enhancers ^28,31,34,37,38^. PCR amplicon Next Generation Sequencing (NGS), which gives the sequence of individual DNA molecules, can be used to define the exact alteration and frequency of sequence variants ^33,34,38^. The only limitation of NGS is cost and turnaround time. Beyond the F0 generation, identifying F1, F2 or beyond that are heterozygous for DNA alterations can be done by Sanger sequencing of PCR products but this approach is very costly. HRMA can define different heterozygous alleles based on the curve shape; however, on first discovery, HRMA will need to be followed by Sanger sequencing to confirm sequence correlation with HRMA curve ^11,33^. However, not all labs have HRMA capabilities. As above allele specific PCR and/or allele specific restriction enzyme digestion is ideal if one can develop a highly stringent PCR assay. NGS would also be feasible, but it is costly, barring a barcoding method to pool individual animal PCRs. The difficulties in exact variant detection add to the challenges in generating an animal carrying the exact desired patient variant. We determined PCR amplicon Next Generation Sequencing (NGS) evaluation of HDR efficiency at the F0 embryo level is best for determining the ideal guide and oligo combination. Further NGS evaluation of DNA from progeny from F0s (germline derived), not F0 biopsy DNA, is essential to identify germline transmitting founders. Surprisingly we find that most founders exhibit a *jackpot* effect in the germ line but not in the somatic tissue. We found NGS superior to using ICE (Inference of CRISPR Edits) for determining HDR frequency. When applicable, allelic-specific PCR or allelic specific restriction digestion can be used to genotype mutation carrying F1 generation animals, however we demonstrated that false positives occur. Further, we successfully used high resolution melting curve analysis (HRMA) to differentiate and identify F1 animals with patient variants.

We have successfully generated 12 zebrafish mutant alleles carrying a single amino acid change. We have applied CRISPR/Cas9 directed oligo homology directed repair (HDR) along with next generation sequencing (NGS) evaluation of HDR rate in F0 embryos and in the identification of high HDR germline transmitting F0 founders. We did not find a consistent length, orientation or symmetry of successful oligos for HDR. We demonstrated the successful use of high-resolution melting curve analysis (HRMA) to detect heterozygous F1 animals and demonstrate the use of allelic specific PCR and allelic specific restriction enzyme digestion as an alternative to HRMA. Altogether we established a highly successful pipeline for the generation of precision zebrafish models.

## RESULTS

### Define analogous patient variant in zebrafish

The first step in the UAB CPAM zebrafish patient variant pipeline (Fig 1) is to determine if that exact amino acid is conserved in zebrafish and if there are one or two orthologs in which to make the variant. The first 7 accepted zebrafish CPAM cases are NF1 R1276Q (RQ), NF1 G484R (GR), VMA21 G55V (GV), SPOP D144N (DN), SGO1 K23E (KE), Pex10 H310D (HD), FKRP C318Y (CY). Amongst these, only NF1 has two orthologs, *nf1a* and *nf1b*, however previous studies have defined the *nf1a* as the predominant protein present in zebrafish ^44^. For NF1 R1276Q (c3827G>A) the amino acids are identical in this region (note R1276 is R1191 in zebrafish), and the wildtype G (c3572G) is conserved (Fig 1B). However, changing only the G to A in zebrafish would code for Lysine not Asparagine, therefore for the mutant allele we will change AG to CA mimicking human mutant CAA codon. For NF1 G484R (2542G>A) the amino acid sequence is identical around the variant site (zebrafish G763) and the G (c2287) nucleotide is also conserved (Fig 1C). To model this in zebrafish we designed the same G>A variant which results in a G>R change. For VMA21 G55V (c164G>T) this amino acid in zebrafish (A58) is different, but the nearby amino acids are conserved. Therefore, we designed the patient C>T variant which will result in an A>V change (Fig 1D). For SPOP D144N (c430G>A) the amino acid sequence is identical around the variant site (zebrafish D144) and the G (c430) nucleotide is also conserved (Fig 1E). To model this in zebrafish we designed the same G>A variant which will generate that patient D>N change. For SGO1 K23E (c67A>G) the amino acid sequence is highly conserved around the variant site (zebrafish K19) and the A (c55) nucleotide is also conserved (Fig 1F). To model this in zebrafish we designed the same A>G variant which will generate the patient K>E change. For Pex10 (c928C>G) the amino acid sequence is highly conserved around the variant site (zebrafish H282) and the C (c844) nucleotide is also conserved (Fig 1G). To model this in zebrafish we designed the same C>G variant which will generate the patient H>D change. For FKRP C318Y (c953G>A) the amino acid sequence is highly conserved around the variant site (zebrafish C355) and the C (c1064) nucleotide is also conserved (Fig 1H). To model this in zebrafish we designed the same C>G variant which generated the patient C>Y change.

**Figure 1:**
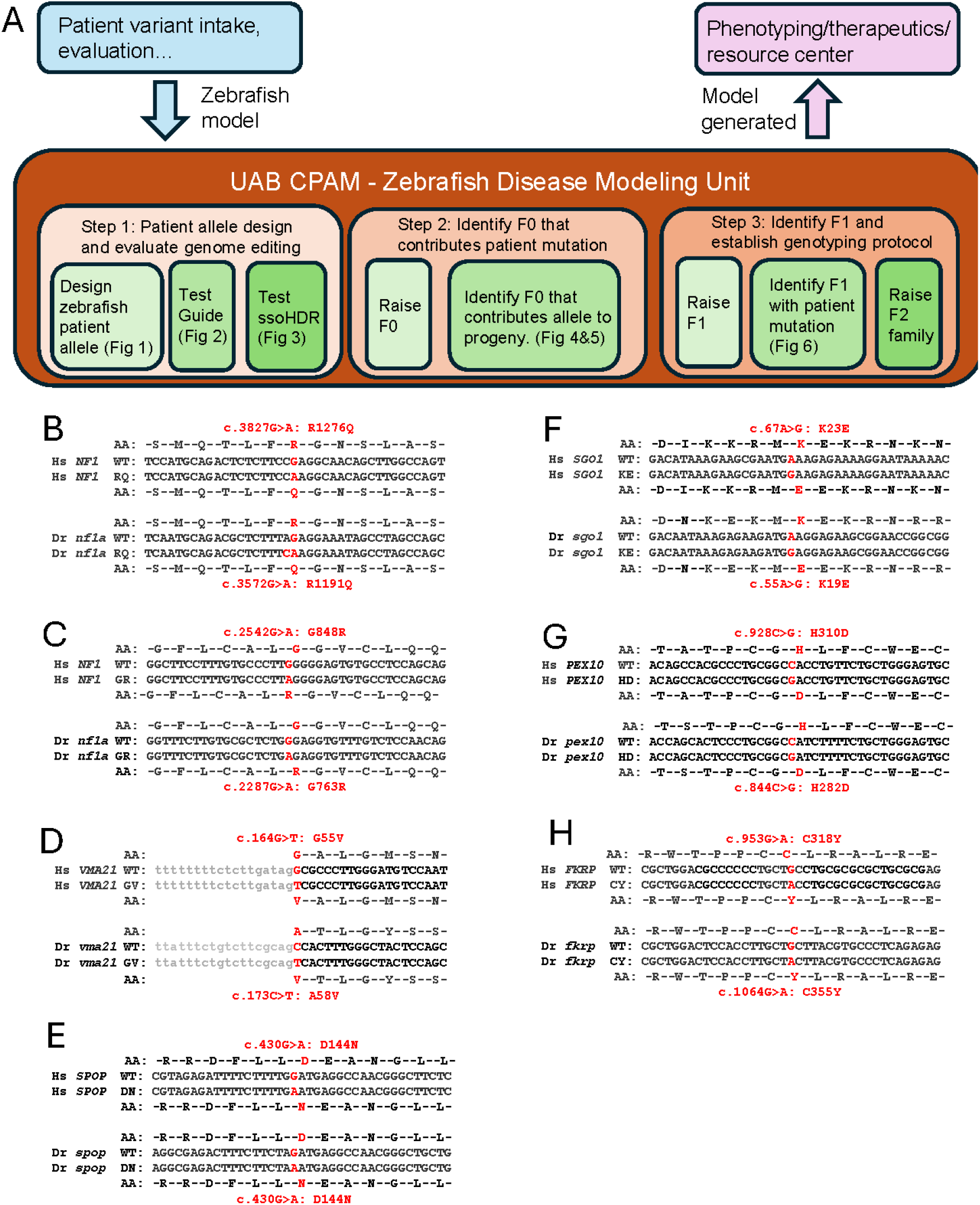
CPAM zebrafish patient variant platform and projects. A) Flow diagram of the CPAM evaluation and zebrafish model production pipeline. B-H) Human wildtype and patient nucleotide sequence and amino acid sequence near the patient variant. This is then aligned with the zebrafish wildtype sequence and potential patient mimetic alteration. Patient change sites are denoted in red font. Exonic sequence is in uppercase, while intronic sequence is in lower case.

### Guide selection and evaluation

The next step in the zebrafish modeling pipeline is to identify potential guides in the vicinity of the codon change based on the NGG PAM (Fig 2A, S1). We have been utilizing the IDT guide design program (https://www.idtdna.com/site/order/designtool/index/CRISPR_CUSTOM) to evaluate the potential guide for on-target and off-target scores (Fig 2B). To evaluate guide directed Cas9 cleavage, we inject the RNP mixture into zebrafish eggs, generate gDNA from 5dpf embryos, and analyze for cutting. Initially our primary assay was HRM analysis (Fig 2B&C, S1), however we have tested ICE and NGS diagnostic approaches as well (Fig 2B&D). Note HRMA is purely qualitative and subjective to users’ discretion, while NGS provides the actual sequence of alleles, ICE predicts the alleles based on the chromatogram. With that, ICE can be influenced by the quality of the chromatogram and “noise” of the run. Our results indicate that while these different methods equate to different empirical numbers the general trend is consistent amongst the techniques (Fig 2B&D; S2). For example, in the *NF1*A R1276Q project, we found that both methods identify the same indels (Δ2(−TT), Δ4(−TCTT), Δ1(−T), Δ11(−AGACGCTCTTT), and a +1(+T) for guide 1; Fig 2D). For labs with HRMA capabilities this is the cheapest and fastest assay, while for most non-HRMA labs ICE analysis will be most effective. Traditionally we have selected the guide with the best on-target and off-target scores, however the need to identify a guide near the variant site often results in lower quality scores. For example, for a *NF1* G848R project, guide 3 had an IDT on-target score of 16, which is low, but based on HRMA (and subsequently NGS and ICE) it was able to cleave at the target site sufficiently (Fig 2B). If cleavage is sufficient among multiple guides tested, we utilized the guide that cleaves closest to the variant site. To evaluate how predictive the scores are with cleavage efficiency, we compared multiple guides with varying on-target scores (Fig 2B). Unfortunately, we have not found the on-target scores to be predictive of cleavage efficiency, suggesting that selecting guides based on location relative to the variant site is the best practice and then evaluate true cleavage efficiency.

**Figure 2:**
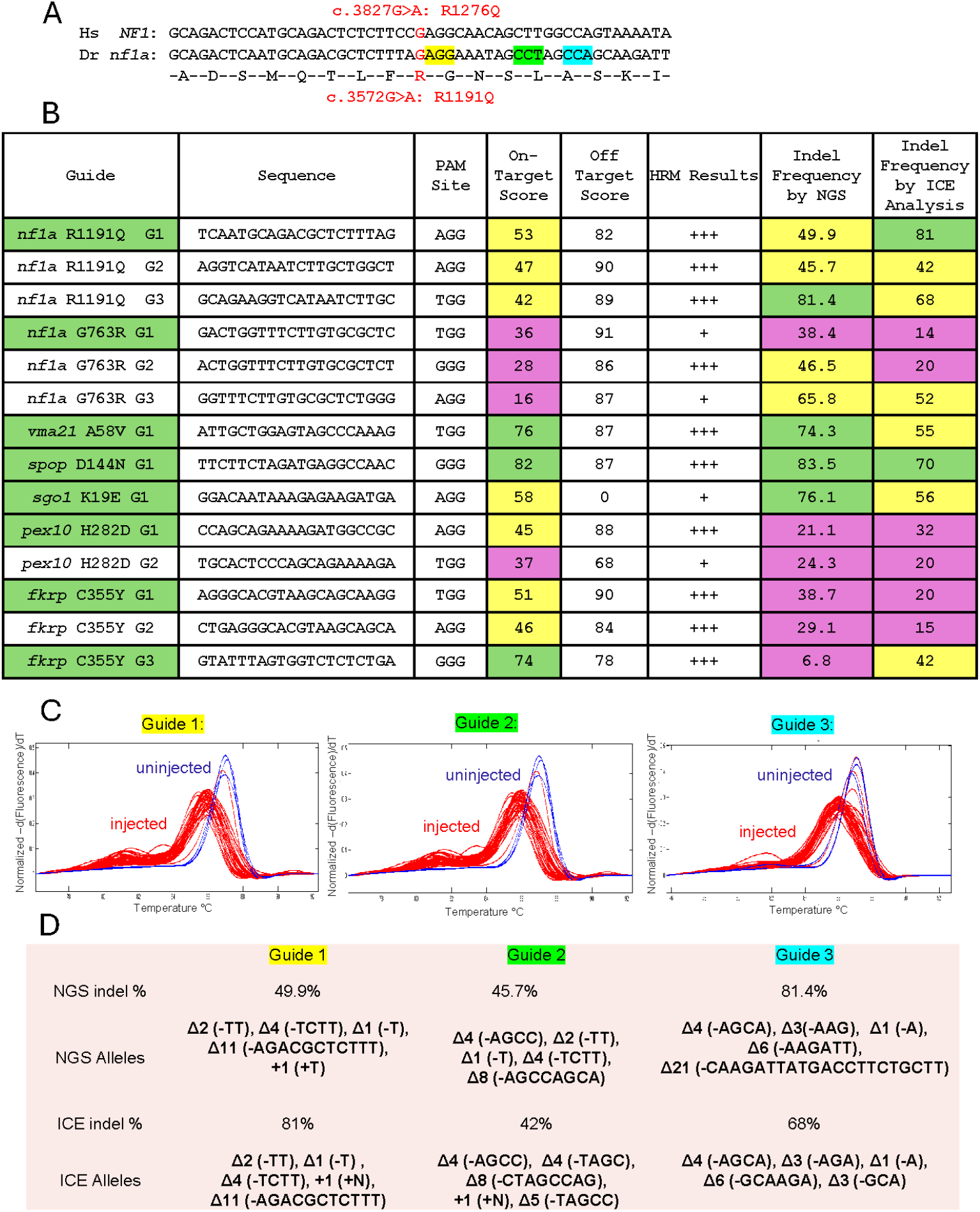
Guide design and evaluation. A) As an example of guide selection, three potential PAM sites for *nf1a* RQ project are depicted. B) Table depicting guides tested, IDT on-target and off-target scores, along with HRMA, NGS and ICE results. Results are color coded to depict strength of result, with green being ideal, yellow being moderate, and magenta being poor. Guide names highlighted in green were used to make the zebrafish line. C) HRMA based evaluation of the guides result in altered HRMA curves (red curves) relative to uninjected (blue curves) indicating guide directed indels generated in the gDNA. D) Evaluation of guide efficiency and identified indels (in order of frequency) by NGS and ICE analysis.

### Evaluating HDR efficiency

To generate the desired patient variant, we inject the defined RNP above along with a single stranded oligo repair template. Studies suggest different sizes and orientations of the oligo can influence HDR, therefore we designed symmetrical long (120nt; 60:60) and shorter oligos (72nt; 36:36) in the PAM strand and reverse complement orientation for the non-PAM strand. We also included asymmetrical oligos (90:36 and 36:90) in the PAM strand and reverse complement (RC) orientation (Fig 3A). In some cases, we designed additional silent mutations to reduce Cas9 recutting (for *sgo1* and *fkrp*, the silent mutation will destroy the PAM site), reduce recombination between the patient change and the cleavage site (for *spop* there are 3 silent mutations; for *pex10* the 2 silent mutations), or humanize the sequence (for *vma21* there are 3 silent changes to humanize the exon sequence and 5 human intron changes) (Fig 3B). To evaluate HDR efficiency, we collect gDNA from pooled (n=∼100) 5dpf injected embryos and perform NGS of the PCR product across the HDR site. For NF1a RQ, we evaluated a template with just the patient mimetic AG>CA change. In the case of the AG>CA template (0 additional changes) the RC 36:96 worked best with an 8% HDR rate, and the 90:36 had the next best with ∼4.2% (Fig 3C). However for the NF1 GR project with 0 additional changes, the 90:36 oligo (1.6%) was best; for VMA21 with 3 additional changes the RC36:96 oligo (3.9%) was best; for *spop* the RC36:90 oligo with 3 additional changes (3.9%) was best; for SGO1 the RC60:60 oligo with 1 additional change (4.1%) was best; for *pex10* the 90:36 oligo with 2 additional changes (1.5%) was best; and for FKRP the RC36:90 oligo with 1 additional change was best (Fig 3C).

**Figure 3:**
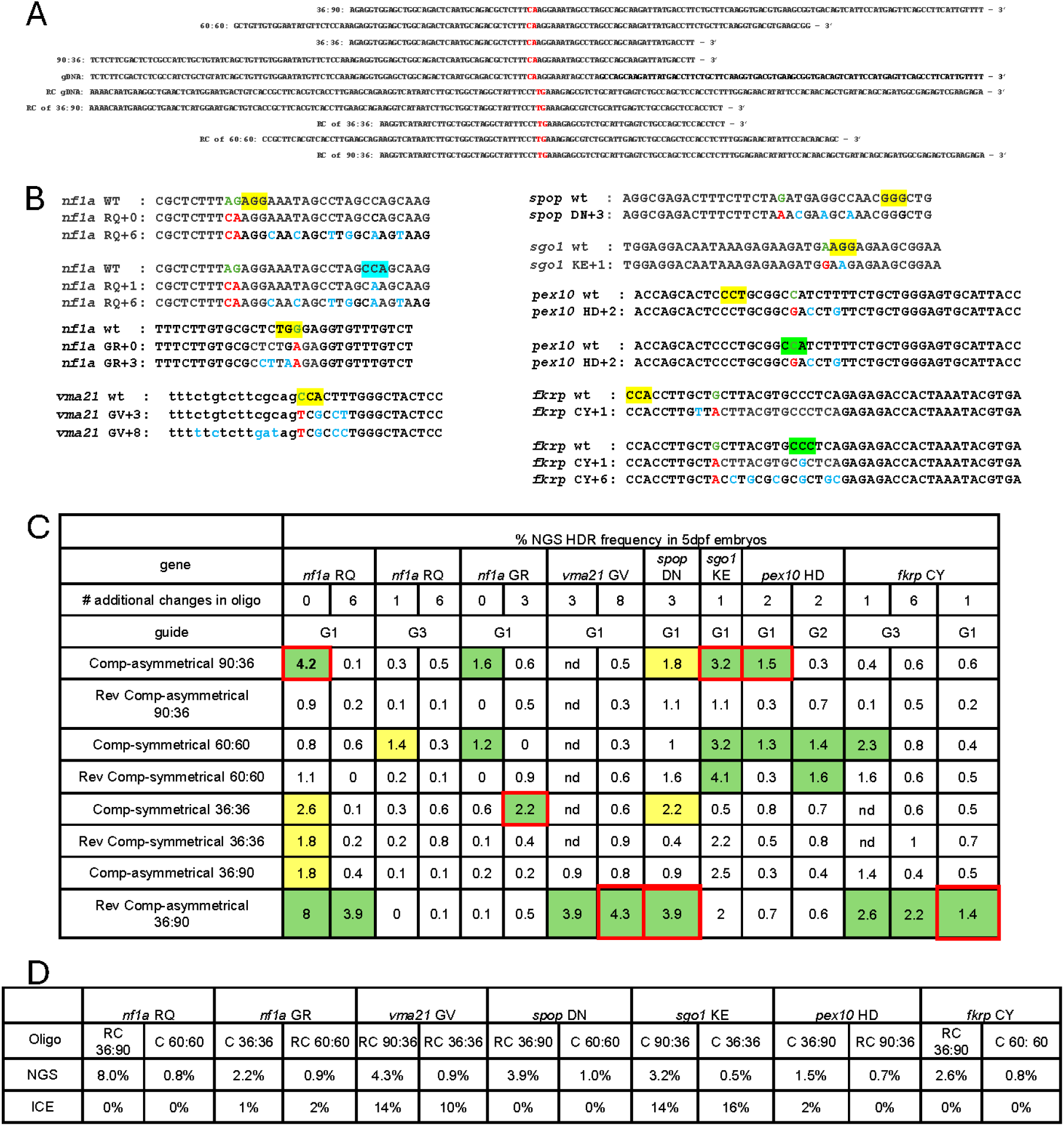
Single stranded oligo design and HDR evaluation. A) Depicts the patient mimetic and mimetic plus additional silent mutations that were used in the oligo design for all the projects. The green font depicts the wildtype sequence, and red font depicts the patient mimetic change. The blue font depicts silent alterations. The yellow, green, or blue highlight denotes the PAM sequence of the guide (G1, G2, or G3) to be used. C) Table summarizing the HDR rates of different oligos using different guides, and with additional alterations. The red border defines combination used to obtain germline transmission of the desired allele; green highlight depicts strong HDR rates, while yellow highlight depicts moderate HDR rates. D) Table comparing HDR rates determined by NGS verse ICE.

Unfortunately, these data did not define an ideal orientation/complement and suggest with each project a different orientation/complement is best and must be determined empirically. That said, if one had to reduce the oligos tested, amongst these cases using both a 90:36 and a RC 36:90 oligo in most projects would generate a sufficient HDR rate. Efficiency of HDR was also affected by the CRISPR guide site location (i.e., closer to the variant site worked better). For *nf1a* RQ, several different oligos worked well with guide 1 (6 different oligos had an HDR rate over 1%; cleavage site 1 bp from variant) but only 1 oligo had an HDR frequency over 1% with guide 3 (cleavage site 20bp from variant). For *fkrp* CY, HDR frequencies initially indicated that several oligos would work with guide 3 (cleavage site 15bp from variant), but we were unable to find a good F0 with this guide/oligo combination but were successful in finding a good F0 with guide 1 (cleavage site 3 bp from variant). However, with the *pex10* HD project, the 2 different guides (cleavage site 2 or 4 bp from variant) worked equally well with the different oligos (guide location was not as critical in this case). We also evaluated the impact of the patient mimetic variant/s and the addition of silent mutations that can help with diagnostics (Fig 3B). In many cases, creating additional silent changes will allow for allelic specific PCR or allelic restriction enzyme sites. However, these silent changes can also reduce homologous recombination. For *nf1a* RQ, we tested a template with the AG>CA and AG>CA plus 6 additional silent changes.

When using the template with 6 additional mutations in all cases the HDR rate was lower, with the RC 36:96 being the best at 3.9% (compared to 8% with GA>CA only; Fig 3C). In the case of *nf1a* GR the oligo orientation preference changed, and the percentage was slightly higher (36:36 with 2.2% compared to 90:36 with 1.6%; Fig 3C). However, in the case of VMA21 and FKRP the same oligo orientation was best and the HDR% did not change much (Fig 3C).

Together this data suggests that extra silent mutations can reduce efficiencies, however in many cases this is tolerable considering the later F1 and beyond diagnostic benefit (allele specific PCR or allele specific restriction enzyme site). After the best oligo plus RNP analysis, we raised the 2 F0 clutches with the highest HDR rate. While NGS is ideal for identifying the exact indel or HDR sequence generated in the embryos, it is costly and if outsourced can require approximately a week to month turnaround time. We evaluated ICE as a cheaper and quicker alternative for determining HDR efficiency. Note while HRMA is fast and cheap it is difficult to differentiate indels verses HDR rates in the F0 population by this method. To evaluate ICE, we compared HDR efficiency with NGS verses ICE of the same PCR product (Fig 3D). Surprisingly we found the HDR rates to be quite disparate between these two methods, suggesting NGS is ideal for evaluation of best oligo use (Fig 3D).

### Evaluation of germline transmission of patient variants in F0 adult animal

The next step in the process is to identify F0 animals that transmit the patient allele through the germline. Our overall approach is to breed adult F0 to the wildtype AB strain, generate gDNA from a pool (n=100) of 5 dpf embryos, PCR across the variant and evaluate each F0 progeny pool by PCR amplicon NGS. For the NF1a RQ project we screened progeny from 12 F0s (derived from the 90:36 oligo injections) and found one F0 that contributed the desired variant at a 15.7% frequency (Fig 4A). It was surprising that while the F0 embryo screen had an average 4.2% HDR, the majority of the F0s gave less than 1% contribution suggesting possibly an individual “jackpot” effect. We continued to use this approach to evaluate the other projects, and found that for the *nf1a* GR 1 of 35 F0 had a 22.7%, for the VMA21 project 2 of 16 had a 19.4% and 14%, for the *spop* project 1 of 16 had 21.3%, for the *sgo1* project 1 of 11 had a 18.3%, for the *pex10* project 1 of 23 had a 6.1%, and for FKRP 0 of 40 had germline contribution (Fig 4A). For the *fkrp* case we decided to redo the targeting with a different guide/oligo and were able to obtain the desired variant in 1 of 17 F0 screened. Potentially having the cut site closer to the variant (14nt away vs 4nt away) in the second approach helped in targeting, although the HDR rate in the F0 embryos was higher (2.6 vs 1.4) with the first guide/oligo combination. Together these data indicate that there are individual F0’s with higher HDR rates of germline contribution than HDR rates in the pooled F0 embryos (i.e. *nf1a GR* germline F0#26 at 22.7% versus pooled F0 at 2.2%), supporting an individual F0 “Jackpot” effect.

**Figure 4:**
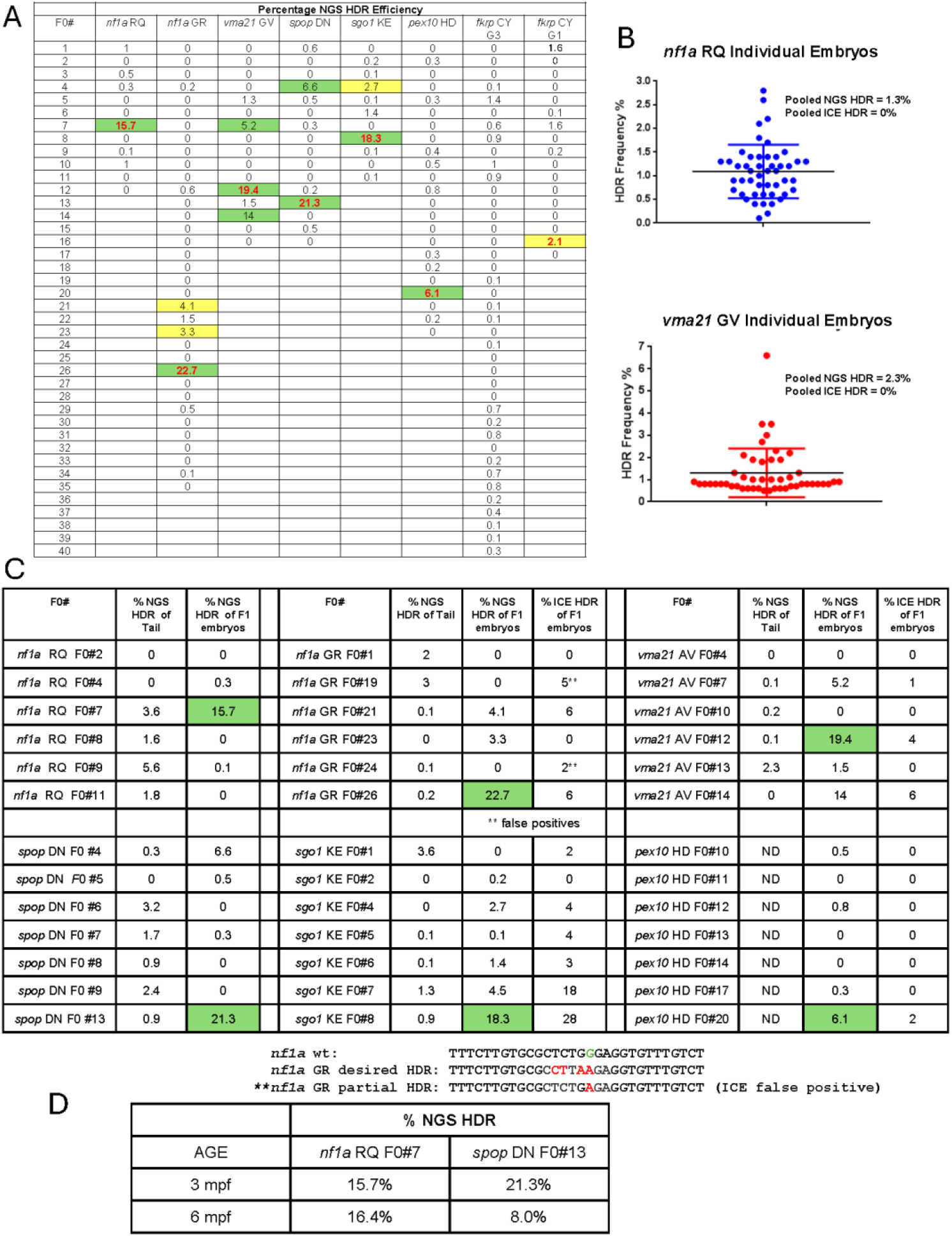
Germline jackpot allele detection in founders. A) Table depicting how many F0 founders were analyzed and the NGS derived HDR rate in pooled (n=100) 5dpf F1 embryos. Animals used to identify adult F1 are denoted with red font. Green highlight denotes founders that give >5% HDR; while yellow denotes >2% HDR. B) NGS based HDR frequency evaluation of individual oligo/RNP injected F0 5dpf embryos. C) Table comparing HDR rates from F0 tail biopsy gDNA and pooled 5dpf F1 progeny gDNA; as well as HDR rates of pooled 5dpf F1 progeny gDNA defined by ICE analysis. Green highlights denote the highest HDR rates observed and the F0 from which the desired F1 was obtained. ** denotes false positives determined by ICE analysis (have only one of the 4 desired HDR alterations but were scored as HDR positive by ICE. D) Evaluation to determine if the germline HDR rate is maintained with age or transient germline populations.

To determine if the jackpot effect exists amongst individual F0 injected embryos, we isolated genomic DNA from 5 dpf individual injected embryos and analyzed these individuals for HDR rates by NGS. For the *nf1a* RQ project (F0 with 15.7%) and *vma21* GV project (F0 with 19.4%) we identified 0 of 48 and 1 (7%) of 48 embryos respectively with a HDR rate above 4% (Fig 4B). This indicated that the F0 germline jackpot effect is not derived from a unique high HDR individual embryo. To further evaluate if the jackpot effect was in the germline only of F0 adults, we analyzed tail gDNA versus F1 progeny gDNA. We tested F0 adult tail gDNA for the *nf1a* RQ, *nf1a* GR, *vma21* GV, *spop* DN, *sgo1* KE, and the *pex10* HD projects. In all cases the adult individual that had high germline contribution did not have high HDR tail DNA contribution (*nf1a* RQ: 15.7 vs 3.6; *nf1a* GR: 22.7 vs 0.2; *vma21* GV: 19.4 vs 0.1, *spop* DN: 21.3 vs 0.9, and *sgo1* KE: 18.3 vs 0.9; Fig 4C). In addition, animals with high HDR in tail DNA did not have high HDR in the germline (*nf1a* RQ: 5.6 vs 0.1; *nf1a* GR: 3 vs 0; *vma21* GV: 2.3 vs 1.5, *spop* DN: 3.2 vs 0, and *sgo1* KE: 3.6 vs 0; Fig 4C). To determine if germline contribution was a transient contribution, we bred the *nf1a* RQ F0#7 and *spop* DN F0#13 again at 6 months and found high germline contribution to be maintained (15.7 vs 16.4 and 21.3 vs 8), but in the case of *spop* DN the frequency was reduced slightly suggesting the number maybe dynamic (Fig 4D). Together this discourages identification of good F0 based on tail DNA and requires testing F1 progeny. This also suggests that HDR events in the primordial germ cells are important, but hard to predict outside of breeding.

To evaluate if ICE could be used in place of NGS, as a quicker alternative, to identify high contribution F0, we split the PCR of gDNA between ICE analysis and NGS. For *nf1a* GR, *vma21* GV, *sgo1* KE, and the *pex10* HD projects while the empirical numbers were different, ICE analysis was able to ID the higher HDR F0 consistent with NGS analysis (22.7% vs 6%; 19.4% vs 4%, 18.3% vs 28%, and 6.1% vs 2%; Fig 4C). However, for some F0s the ICE result suggests high germ line contribution, while NGS would not (*nf1a* GR: 5 vs 0, and *sgo1* KE: 18 vs 4.5; Fig 4C). Considering the NGS is actual reads we interpret the ICE results as false predictions. In the case of *nf1a* GR F0#19 and #24, this was also apparent by the predicted ICE sequence in which only 1 of 4 designed nucleotide changes was called positive (Fig S3A), whereas in NGS only the exact desired seq is called positive (Fig 4C). In another case (*fkrp*) in one F0 there was an additional nucleotide change identified by NGS not in the designed sequence that was called positive by ICE (Fig S3B). Also, some positives called by ICE were not positive by subsequent NGS analysis (Fig S3C). Together this suggests that ICE could be used as a cheaper and quicker approach but does have limitations with calling of false positives. Potentially ICE could be used as a primary screen of pooled F1 progeny, and then NGS only Ice positive pools. Or individual (not pooled) F1 embryo from ICE positive F0 can be PCR amplified, and Sanger sequencing could be used to confirm the F1s that carry the desired patient variant.

Allele specific PCR is an alternative approach to identify good F0. Therefore, we designed a forward primer that is specific to the mutant sequence (Fig 5A). For allele specific PCR to be useful at the F0 stage, the PCR needs to be specific to the patient allele and sensitive to low percentage chimerism. Toward the former, we test specificity of allelic specific PCR on 2 wildtype gDNA samples (Fig 5b). The *nf1a* RQ, *sgo1* KE, and *fkrp* CY primer set amplified a product on the wildtype DNA excluding them for use for allelic specific PCR. This is likely a consequence of only having 2 nucleotide changes between the wildtype and patient allele. *Spop* DN primer set also produced a weak product, establishing concerns that it will have false positives with this primer set. *Nf1a* GR, *vma21* AV, and *pex10* HD primer sets did not amplify a product from wildtype gDNA. This specificity likely results from the additional silent mutations in these sequences (*nf1a* GR with 4 total changes, *vma21* with 9 total changes, and *pex10* with 3 total changes). To test patient allele specificity, we next PCR amplified gDNA from sequence identified patient variant containing F1 gDNA. In each case, the allelic specific PCR was able to amplify a product in gDNA from the F1 animals but not the wildtype animals (Fig 5C, D, & E lanes 2 and 3 vs 4). To test the sensitivity of this approach, we used these allele specific PCR primers on gDNA from the same pooled F1 (n=100) progeny of F0 for which we already know the NGS HDR rate. For *nf1a* GR we could detect a PCR product in F0s with 3.3%, 4.1%, or 22% HDR, but not in F0s with rates below 1.5% (Fig 5C). For *vma21* we could detect positives in F0s with 14% or 19.4% HDR but did not in any F0 with a rate below 5.2% (Fig 5D). For *pex10* we could detect positives in the F0 with 6.1% but did not in the F0 with a 0.3% HDR rate (Fig 5E). Together this data indicates that allelic specific PCR is a reasonable alternative for identification of good F0 animals, however, this may be restricted to detecting only high-level chimeric animals, and benefit from incorporation of extra silent mutations in allele design.

**Figure 5:**
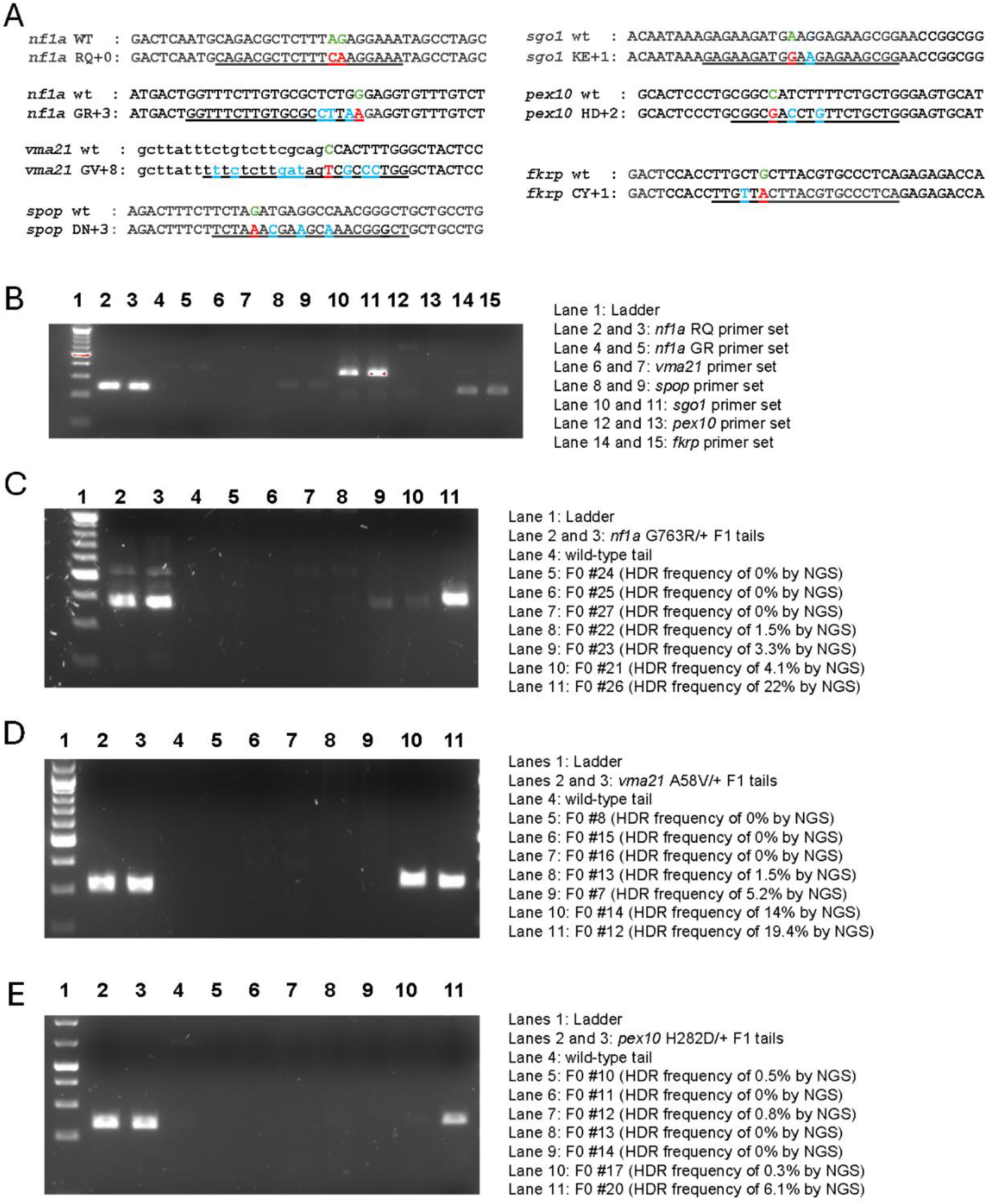
Using allelic specific PCR to identify germline transmitting founder animals. A) Allele specific PCR forward primer (underlined) design for each project. B) Use of each allele specific primer set with wildtype gDNA to determine if mutant specific primer sets amplify a product off of wildtype gDNA. Two wildtype tail biopsies derived gDNA samples are used per primer set. C) Agarose gel of PCR products amplified from *nf1a* GR F1 heterozygous and wildtype control tail gDNA and pooled 5dpf F1 progeny gDNA from NGS HDR rate defined F0 founders. 359 bp band denotes the presence of the HDR allele. D) Agarose gel of PCR products amplified from *vma21* AV F1 heterozygous and wildtype control tail gDNA and pooled 5dpf F1 progeny gDNA from NGS HDR rate defined F0 founders. 303 bp band demotes the presence of the HDR allele. E) Agarose gel of PCR products amplified from *pex10* HD F1 heterozygous and wildtype control tail gDNA and pooled 5dpf F1 progeny gDNA from NGS HDR rate defined F0 founders. 234 bp band denotes the presence of the HDR allele.

### Detection of F1 animals carrying the patient variant

The next step in the pipeline is the identification of F1 adults carrying the desired patient variant. There are many approaches to genotype animals ranging from HRMA being the cheapest and fastest to individual animal Sanger sequencing being the costliest per sequencing reaction. The choice of method depends on individual lab capabilities, but also how many animals will need to be screened to get a positive. The highest germline transition rate was 22.7 (*nf1a* GR #26) which should produce HDR positives in 1 of 5 F1 adults. The lowest germ line transmission rate was 2.1% (*fkrp* CY F0#16) which we would expect 1 in 47 F1 adults. HRMA approaches do not differentiate the actual alleles; they just define if a F1 has a sequence difference. However, in many cases the shape of the HRMA curve can differentiate alleles. For example, based on the NGS data from pooled gDNA of F1 progeny from *nf1a* RQ F0#7 should also contribute the following indels Δ17 (9.9%), Δ2 (3%), +2 (3.5%) alleles. For *nf1a* GR we were able to differentiate F1 with 3 different alleles, and then through Sanger sequencing we defined them as GR/+ (16%), Δ17/+ (8%), Δ2/+ (16%), and +2/+ (8%) (Fig 6A). For *fkrp* CY F0#16 the pooled F1 embryo data predicted indels Δ17#1 (33.8%), Δ17#2 (4.7%), Δ1 (3.8%), and Δ4 (3.4%). Using HRMA we were able to differentiate F1 with 5 different alleles, and then through Sanger sequencing we defined them as CY/+ (6.4%), Δ17#1 (7.4%), Δ17#2 (4.3%), Δ1 (8.5%), and Δ4 (7.4%) (Fig 6B). For *nf1a* RQ, *vma21* AV, *spop* DN projects we were also able to use HRMA to differentiate F1 animals based on HRMA curves (Fig S4). Note for *sgo1* KE we could differentiate F1 that carry an altered allele relative to wild type but could not differentiate the patient allele versus indels using HRMA. For this project we used sanger sequencing of HRM positive F1 to identify the patient variant F1 carrier and bred this to AB to generate F2 KE carriers with the patient variant. These could be genotyped by HRMA.

**Figure 6:**
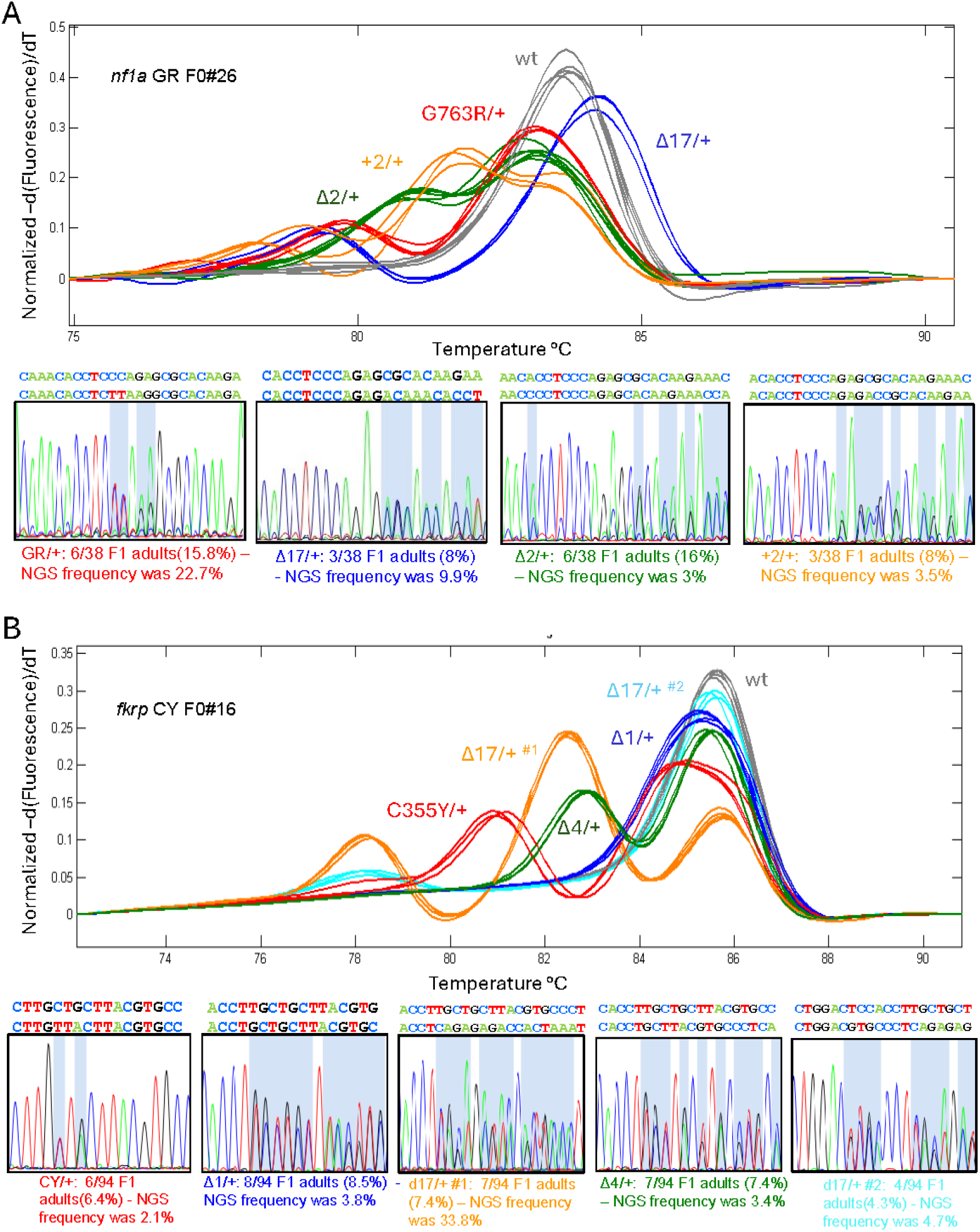
HRMA detection of adult F1 patient carrying mutations. A) Example of HRM analysis of F1 adult tail biopsies from *nf1a* GR F0#26 (22% HDR rate in pooled F1 embryos). Different curves and associated allelic sequences are denoted by colored curves. Below are Sanger sequencing chromatograms of gDNA from F1 animals with different curves; along with the frequency of the allele in 38 F1 analyzed and the pooled F1 embryos NGS data. B) Example of HRM analysis of F1 adult tail biopsies from *fkrp* CY F0#16 (2.1% HDR rate in F1 pooled embryos). Different curves and associated allelic sequence is denoted by colored curves. Below are Sanger sequencing chromatograms of gDNA from F1 animals with different curves; along with the frequency of the allele in 94 F1 analyzed and the pooled F1 embryos NGS data.

HRM analysis is not available to all researchers, so we also evaluated allelic specific PCR and restriction digest allelic PCR for those designs applicable. We wanted to test how well these primer sets can specifically detect F1 animals with the specific variant versus indels that come from the same F0. For *nf1a* GR, *vma21* AV, *spop* DN, and *pex10* HD we were able to amplify a PCR product on the patient allele carrying F1 (Fig 7A-D). However, for *nf1a* GR specific PCR and *spop* DN specific PCR we observed a PCR product in the *nf1a* +17/+ and *spop* +20/+ (1 of 2 samples) lines (Fig 7A&B). In the case of +17, this new inserted sequence is a perfect match for the allele specific primer (Fig 7A). In the case of 1 of 2 Spop +20/+ samples we think this is not specific to this allele, but more stochastic and reflects the ability to weakly amplify the wildtype allele (Fig 7B). Note amplification of the HDR allele is stronger. Together this suggests allelic specific primers can be used to ID F1, but sequencing of these positives is required to validate they are the correct sequence. In the case of *spop* DN, the patient allele destroys an XbaI restriction site (Fig 7E). Therefore, we wanted to determine if digestion of the PCR product could be used to detect patient allele F1 animals. We were able to demonstrate that this approach can effectively identify DN heterozygous F1 animals, but not wildtype or Δ8 or +20 heterozygous animals (Fig 7E). Of note, while in this case the deletion allele did not destroy the enzyme site, often if the cleavage site overlaps with the enzyme site the indel will destroy the enzyme site. If this approach is to be used, making the patient allele have a new restriction site would be more specific to the patient allele.

**Figure 7:**
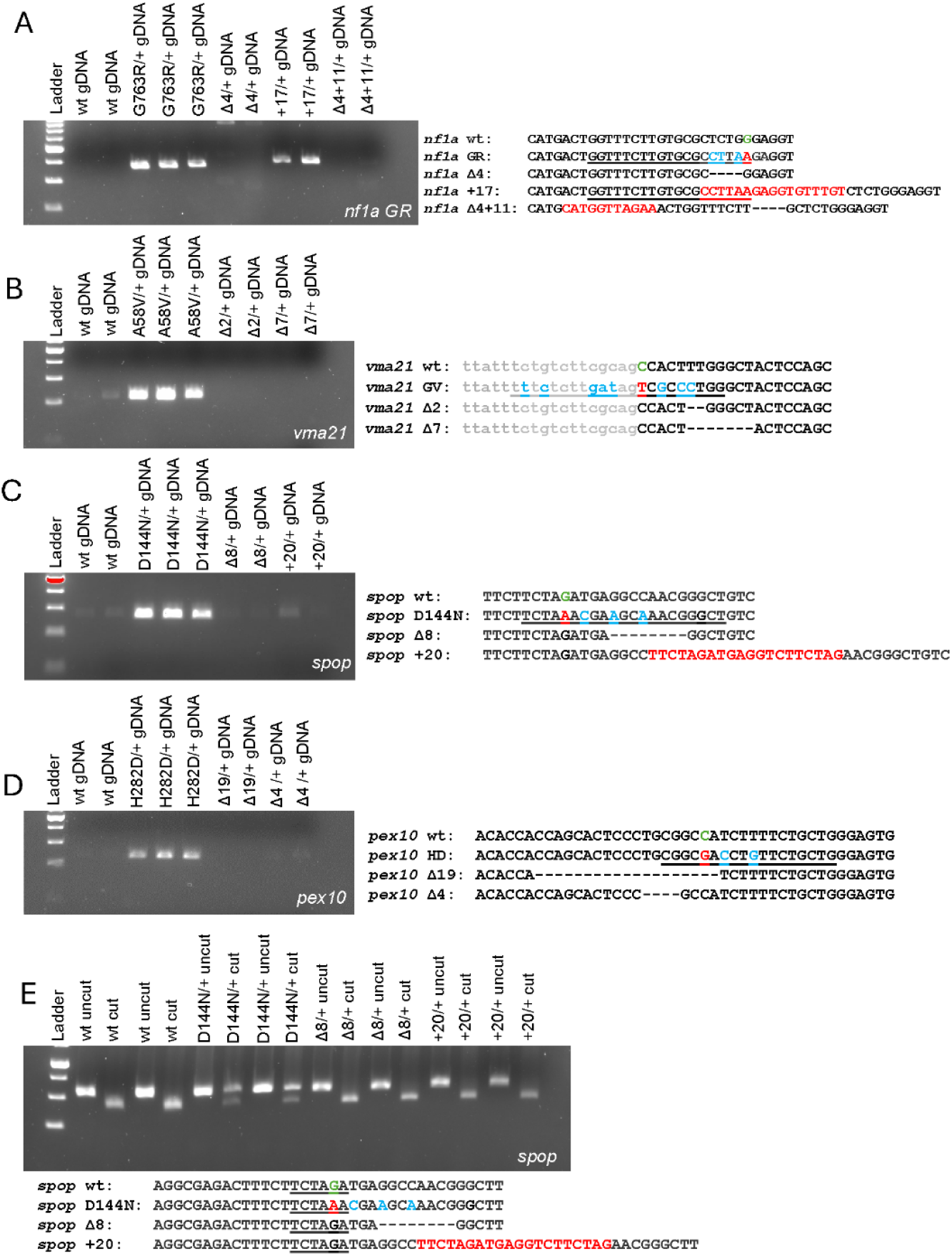
Allelic specific PCR and allelic specific restriction digestion of F1 patient variant carrying animals. A) Agarose gel of PCR products using *nf1a* GR allele specific PCR of F1 heterozygous patient variant and heterozygous indels. 359 bp band denotes the presence of the HDR specific allele. B) Agarose gel of PCR products using *vma21* AV allele specific PCR of F1 heterozygous patient variant and heterozygous indels. 303 bp band demotes the presence of the HDR specific allele. C) Agarose gel of PCR products using *spop* DN allele specific PCR of F1 heterozygous patient variant and heterozygous indels. 274 bp band denotes the presence of the HDR specific allele. D) Agarose gel of PCR products using *pex10* HD allele specific PCR of F1 heterozygous patient variant and heterozygous indels. 234 bp band demotes the presence of the HDR specific allele. Sequence of wt, HDR allele, and indel are depicted below. Allele specific primer is denoted by underlined sequence in the HDR allele. E) Agarose gel of restriction enzyme digested *spop* PCR product of F1 heterozygous patient variant and heterozygous indels. 321bp band denotes the uncut product, while 273 and 48 bp bands denote the cut product. Sequence of wt, HDR allele, and indel are depicted below with the XbaI restriction enzyme site underlined in the D144N sequence.

### Second round of zebrafish model generation

To further demonstrate effective zebrafish model generation using this CRISPR HDR platform we generated 4 additional patient alleles (NF1 R681*, NF1 M992del, P53 R175H, and PKD2 L656W) and 1 research initiated allele (*p53* K117R). For all these projects, the amino acid and surrounding sequence is conserved between human and zebrafish (Fig 8A). Human NF1 R681 is *nf1a* R628, NF1 M992 is *nf1a* M907, P53 R175 is *tp53* R144, PKD2 L656 is *pkd2* L593 and *p53* K120 is *tp53* K89 in zebrafish. Efficient guides near the variant site were validated using HRM analysis of injected 5dpf F0 pooled embryos (Fig 8B). For *nf1a* R628, while the patient change is C>T, in zebrafish the analogous change is a A>T change generating the TGA stop codon (Fig 8A). For *nf1a* M907del, in patients the AAT is deleted, therefore we designed the oligo to delete the analogous CAT in zebrafish (Fig 8A). For *p53* R144H, the patient alteration is a G>A in codon CGC, however in zebrafish we had to alter the AGA codon to CAC to mimic the patient variant (Fig 8A). For *pkd2* L593W, the patient variant is a T>G, therefore in zebrafish we designed the same T>G change plus an additional silent G>C variant to discourage Cas9/Guide directed recutting (Fig 8A). For the *p53* K89R, we designed the zebrafish allele to mimic the mouse A>G change ^45^, but added a silent mutation to discourage Cas9/Guide directed recutting (Fig 8A). Effective HDR oligos were identified by NGS of injected 5dpf F0 pooled embryos (Fig 8B). F0s from effective CRISPR HDR oligo combinations were raised and bred to AB to identify germline contributing founder animals. For *nf1a* R681* we screened 5dpf pooled F1 progeny by ICE analysis, followed by individual embryo Sanger sequencing. From this we defined 6 ICE positive founders out of 6 screened. However, when we sequenced individual F1 embryos from 3 of these ICE positive founders only one F0 provided germline F1 progeny by ICE (Fig. S3D). We believe this high false positive rate in the pooled F1 is related to inaccuracy of ICE analysis when germline contributions are low. In the next 4 projects, we used NGS of progeny from F1 founders to identify good founders (Fig 8D). In each case, we were able to identify individual adult F1 carrying the desired variant within as many as 14 founders. In *nf1a* M907del we identified positives F1s in as low as 1 of 4 founders. These further demonstrate the effectiveness of this zebrafish editing platform.

**Figure 8:**
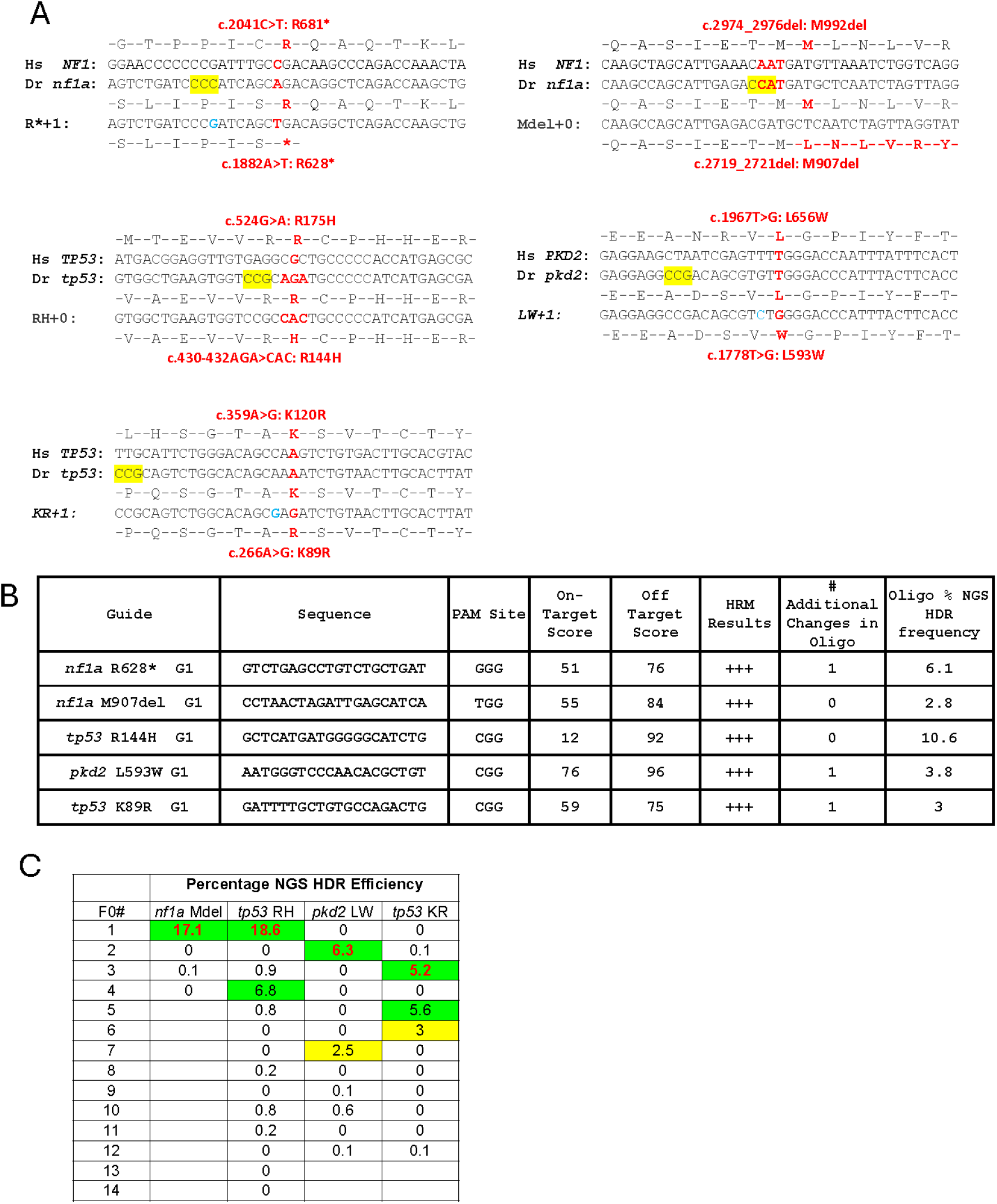
Generation of 5 additional zebrafish models: A) For NF1 R*, NF1 M992del, TP53 R175H, TP53 K120R and PKD2 L656W models, human wildtype nucleotide and amino acid sequence near the patient variants. This is then aligned with the zebrafish wildtype sequence and potential patient mimetic alteration. Patient change sites are denoted in red font. B) summary of guides used, IDT scores, HRM results, and NGS derived HDR frequencies. C) Frequency of positive F0 and the germline transmission frequency for the projects.

## DISCUSSION

Cell based models are easy and fast, however they do not recapitulate the complex cellular interactions, cell fate changes during development and repair, and cell type specific aspects of a disease. These characteristics are often best studied in animal models. Zebrafish is growing in utilization as a model of human disease, initially due to its conserved genome relative to human, short generation time, large brood size (>200 eggs per pair), external fertilization, and transparent embryos. These advantages have expanded to 1) genetic screens for disease phenotypes, 2) ease of transgenesis using tol2 transposases, 3) morpholino or crispant screen for genotype phenotype correlations, 4) large cohorts for cancer survival analysis, 5) the regenerative capacity of zebrafish has gained much attention toward understanding how to regenerate tissues, and 6) zebrafish also serve as an ideal model for *in vivo* chemical screens. The latter has gained much attention in the patient modeling arena, due to the ability to evaluate drug repurposing efforts quickly and efficiently. Together zebrafish models of human disease have become useful not only for pathology but also for therapeutics.

Recent advances in genome sequencing have illuminated many disease-causing genes/variants. The fact that specific variants can propagate specific disease phenotypes often not associated with loss of function (LOF) mutations has warranted a better understanding of the in vivo pathobiology of these variants. To simplify aspects, a patient variant can act as dominant negative (DN), gain of function (GOF), hypomorphic alleles, or affect only a certain protein domain function of the protein. Patient variants can have DN effects in which the mutant protein binds the wildtype protein and inhibits its ability to function, which is most often the case with proteins that homodimerize. Importantly this phenotype will often present itself in a heterozygous individual. A GOF effect usually refers to an activity or phenotype associated with a mutant allele that is not associated with loss of function alone; For example when homozygosity for a missense allele results in a more severe phenotype than homozygous null for a gene. This is often derived from the mutant protein binding to a complex and making the complex non-functional, in essence making all the protein in the complex similar to being null. Some GOF variants can result in an enzymatically active state. While less clear, sometimes GOF variants will have a new activity, such as if the variant allowed the protein to have a new binding partner. GOF phenotypes are usually defined by comparing the homozygous null to the homozygous variant. Hypomorphic alleles usually have less activity. In some cases, in which complete loss of activity is lethal, patients with hypomorphic alleles may live but have disease phenotypes. The easiest way to access if a patient variant is hypomorphic in nature is to compare the phenotype of a homozygous variant to a wildtype and homozygous null. If the homozygous patient variant has a “weaker” phenotype than the null, but more severe than the wildtype, it is likely hypomorphic. In some cases, the homozygous patient variant is equivalent to the homozygous wildtype, in which it is best to sensitize the analysis by breeding the patient variant over a null, to see if this forms a phenotype relative to the wildtype. Understanding genetics using animal models helps direct future molecular research on the pathobiology of these patient variants and can inform on approaches to develop therapies.

In the majority of CPAM projects we have found high homology at the patient variant amino acid in zebrafish. This by itself could suggest the importance of this amino acid for protein function since it is conserved across species. In some cases, like VMA21 the amino acid is different, but our analysis indicates that the alteration occurs near a splice junction disrupting normal RNA processing that is the main source of the patient phenotype. In this case, we felt it was best to humanize some of the intron and the exon junction.

There was a genome duplication and reduction in teleosts, including zebrafish, that in some cases resulted in 2 orthologues. The consequence of this is diverse; in some cases, promoter/enhancer elements have diverged controlling expression of the gene in different domains, while in others the dominance of the genes varies. For nf1, *nf1a* is the more impactful gene, while *nf1b* contributes to overall nf1 function^44^. In another project *ebf3a* null results in a phenotype and *ebf3b* null does not have a phenotype or does not enhance the phenotype in the double null ^23^. In DDX3X, zebrafish *ddx3a* or *ddx3b* null are viable, while double null is early lethal (unpublished/submitted data). For patient modeling, this duplication leads to three decisions regarding modeling: 1) don’t model in zebrafish, 2) model in both orthologs, or 3) model in the more essential/dominant ortholog or ortholog expressed in disease tissue of interest, and study over the null in the other ortholog. We favor the last, due to ease, but could see a situation where the patient variant does not model the disease due to not being expressed in a specific tissue therefore having a dominant negative or gain of function activity in that tissue.

The overall approach to generating patient variants in zebrafish is more similar to generating mutation in cells, than rodents. In rodents, since early development is very slow, taking 24 hours to transition from 1 to 2 cells, and 2 to 4 cells, HDR events can produce animals that are homozygous for the variant. In zebrafish, early cell division takes 20 minutes, making it unlikely the HDR event will happen at the 1 or 2 cell stage. In fact, it is rare to get high chimeras for a specific indel, which have a higher event frequency than HDR. We often observe 2-7 different indels go germline from founders suggesting there are multiple alleles in the germ line. This does allow for the generation of allelic series of indels. The challenge this brings is to define the patient allele in chimeras with indels as well. One key advantage of zebrafish for this is the ability to obtain and raise hundreds of progeny from a single mating. This allows us to obtain rare alleles from animals with low HDR rates. Our HDR data is summarized in Fig S5, which indicates the worst F0 frequency was 4.4% (*pex10* HD; requiring 23 F0’s), excluding the failed first attempt at *fkrp* CY, while the worst germ line transmission rate was ∼5% (*pex10 HD* and *pkd2 LW*; requiring ∼20 F1 animals). The key to obtaining these manageable frequencies is to optimize or preselect for the best HDR cocktail with the highest HDR rate so that identification of the allele is not too cumbersome. Therefore, we find it to be important to evaluate the HDR rate of different cocktails in normal looking F0 injected embryos at 5 dpf. We chose 5 dpf, since this allows for the HDR event to occur, while also excluding any lethality that is associated with biallelic null indels. Some of the variables in these cocktails involved the different orientation and size of oligos. In some cases where we observe high amounts of lethality, we will reduce the guide or Cas9 concentration to lower the prevalence of biallelic indels. Importantly once we find the most efficient HDR cocktail we have been successful in generating all patient variant models. We should note, although we have not observed this, that some patient variants could be heterozygous lethal, in which we could observe the patient variant allele in the 5 dpf F0 embryos but not obtain adult founders carrying this allele. Alternatively, one could get low level chimeras to survive but the F1 progeny does not. This can be determined by assessing if 5 dpf F1s exist with the patient variant but not the adults. In this case one would need these chimeras to access the patient variant phenotype in the F1 generation.

Base editors are an attractive alternative to oligo HDR due to the lack of indels generated. However Base Editors can only be used for A>G (T>C) changes using adenine base editors or C>T (G>A) using Cytosine bases editors. And the desired changes have to be in a restrictive 4 nucleotide window of the guide. Amongst these 12 projects, 5 have A>G or G>A alterations, however none of our tested guides have an editing window that would target the desired nucleotide. That said, newer base editors using PAM-less variants allow for more flexibility in the guide sequence and could overcome this restriction ^46^. Confounding the isolation of the desired variant, most of these have neighboring As or Gs in the editing window likely resulting in multiple bases edited forcing evaluation of many F1s for the desired alteration by sanger sequencing. Further, the base editing approach does not allow for the insertion of tailored silent restriction enzyme mutations.

We observed that somatic HDR rates don’t match the germline HDR rates. In fact, we observed germline transmission rates much higher than we ever see in individual embryos. For example, for *nf1a* RQ the highest individual embryo HDR was 2.7, while the highest germline was 15.7%. This could relate to a limited number of primordial germ cells that will make up the germline. Such that if an HDR event happens early in one or more of these primordial germ cells (PGC) the animal will have a high germline transmission rate. Importantly this makes it hard to predict which founder to use based on somatic tissue analysis and thus required analysis of germline HDR rates. This also highlights that PGC specific genome editing would be ideal for future patient variant genome editing. It would still require assessment of HDR in the F1 progeny of F0 animals but could circumvent lethality often associated with biallelic CRISPR/Cas9 induced indels of essential genes. We envision this might involve the generation of a germ cell specific Cas9 line in which we would inject the guide into or use nanos tagged Cas9 mRNA which will localize to PGCs. One question still unanswered is how efficiently endogenous RNP complexes will be generated in vivo as opposed to when we generate these complexes in vitro and then inject them ready to cleave guide directed sites.

Whenever using CRISPR/Cas9 there are always concerns of off-target alterations. While this concern is extremely relevant to cell lines, animal models can be bred out to the wildtype strain multiple generations to “clean up” the background and exclude off target mutations. In essence the first outcross adds 50% “clean” genomic DNA and then the subsequent crosses result in 75%, 87%, 93%, 96% and so forth. To add to that, the likelihood of breeding two heterozygous carriers for an off-target alteration is low. But this does point to the need for future good phenotyping practices by characterizing the phenotype in at least 3 independent heterozygous pair derived clutches. If the phenotype is identical then it is the patient variant derived phenotype. If there is availability between the clutches, then there may be concern of off-target alterations in the background and additional outcrosses should be performed. Off target genotyping can be used to accelerate selection of animals without alterations, but this is limited to how well the off-target algorithms are. Breeding to wildtype strain multiple generations will remove non-linked genomic alterations.

## METHODS

### Zebrafish Lines and Maintenance

All zebrafish work was performed at the University of Alabama at Birmingham in the Zebrafish Research Facility (ZRF). Adult fish and embryos are maintained as described by Westerfield et al (1995) by the ZRF Animal Resources Program which maintains full AAALAC accreditation and is assured with OLAW. All animal studies have UAB IACUC approval. All zebrafish lines are generated and maintained on the AB strain.

### RNP and RNP/oligo Preparation and Microinjection

Both human and zebrafish genomic and protein sequences were obtained from ensembl.org to locate the exact site of the human patient variant and the site of the desired zebrafish variant and possible crRNA sites. Alt-R crRNA target sites were designed with Integrated DNA Technologies Alt-R CRISPR HDR Design Tool (https://www.idtdna.com/pages/tools/alt-r-crispr-hdr-design-tool). RNP mixtures of Alt-R CRISPR-Cas9 crRNA, tracrRNA (IDT, 1072532) and Alt-R S.p. Cas9 Nuclease V3 (IDT, 1081058) were prepared following manufacturer’s instruction. A 3µM gRNA solution is obtained through diluting 3µl of 100µM crRNA and 3µl of 100µM tracrRNA into 94µl of Nuclease-Free Duplex Buffer, heating at 98°C for 5 min, then cooling to room temperature. 0.5µL Cas9 protein was diluted with Cas9 working buffer (20mM HEPES; 150mM KCl, pH7.5) to yield a working concentration of 0.5µg/µL. The diluted Cas9 protein working solution was mixed 1:1 with the 3 µM gRNA solution and then incubated at 37 °C for 10 min.

Microinjection was performed by injecting 1nL of RNP complex into the yolk of 1-cell stage wild-type zebrafish embryos. RNP complex was freshly prepared and left on ice until microinjection. For generating patient variant models, HDR oligos (Table S1) were designed and ordered (IDT) to include the desired patient and/or silent mutations in different orientations (shorter/longer, asymmetrical/symmetrical and +/-strand) and mixed (1uM) with the RNP, and then 1nl injected into one-cell eggs.

### CRISPR Guide evaluation with HRMA

For indel efficiency evaluation, genomic DNA was extracted from ∼24 5dpf injected embryos and evaluated with HRMA (see below). Genomic DNA was isolated from single embryos that were incubated at 98°C for 10 min in 30µl 25mM NaOH in a 96 well plate; then neutralized with 30µl of 40mM Tris-HCl. PCR reactions contained 1ul of LC Green Plus Melting Dye (Biofire Defense, BCHM-ASY-0005), 1µl of 10x enzyme buffer, 0.2µl of dNTP Mixture (10mM each), 0.3µl of MgCl_2_, 0.3µl of each primer (10µM) (Table S2), 1µl of genomic DNA, 0.05µl of Genscript Taq (E00101), and water up to 10µl. The PCR reaction protocol was 98°C for 30 sec, then 45 cycles of 98°C for 10 sec, 59°C for 20 sec, and 72°C for 15 sec, followed by 95°C for 30 sec and then rapid cooling to 4°C. Primers used for HRMA are designed with Primer3. Following PCR, melting curves were generated and analyzed using the LightScanner instrument (Idaho Technology) over a 65-95°C range to determine if the CRISPR guide cuts at the desired site.

### Determining HDR frequency by Next Generation Sequencing and ICE Analysis

To measure HDR frequencies, genomic DNA was isolated from pooled 5dpf embryos by incubating embryos at 98°C for 20 min in 200µl 25mM NaOH; then neutralized with 200µl of 40mM Tris-HCl. PCR reactions contained 6µl of 5x Phusion HF buffer, 0.6µl of dNTP Mixture, 1.2µl of each primer (10µM) (Table S2), 3µl of genomic DNA, 0.3µl of Phusion Hot Start II DNA Polymerase, and water up to 30µl. The PCR reaction protocol was 98°C for 30 sec, then 45 cycles of 98°C for 10 sec, 57°C for 30 sec, and 72°C for 30 sec, followed by 72°C for 4 minutes and then rapid cooling to 4°C. The products of all first round PCR reactions were then purified with the Promega Wizard SV Gel and PCR Cleanup System (Promega, A9282). We used 800 ng of the purified PCR products for sample indexing with the NEBNext Ultra II DNA Library Prep Kit for Illumina (New England Biolabs E7103L) in preparation for next-generation sequencing (NGS) using Illumina NEBNext Multiplex Oligos indexing primers. The samples were run on the Illumina MiSeq platform using 2 × 150bp paired end reads and HDR efficiency was determined using the CRISPR RGEN Tools Cas-Analyzer software available online at http://www.rgenome.net/cas-analyzer/#!^47^. Several of these same purified PCR products were submitted to Azenta for Sanger Sequencing and analyzed with the Synthego ICE Analysis software (https://ice.synthego.com/#/) to compare HDR efficiency rates detected by each method.

### Identification of Alleles by HRMA and Sanger Sequencing

To identify mutated alleles, DNA is extracted from a tail biopsy from F1 or F2 progenies in each mutant. Tails were collected in 96 well plates and were incubated at 98°C for 20 min in 40µl 25mM NaOH; then neutralized with 40µl of 40mM Tris-HCl. HRMA on the LightScanner instrument is performed to identify the different alleles present and then samples representing the different HRMA curves are sequenced. The DNA fragment flanking the targeting site is PCR amplified using Phusion Hot Start II DNA Polymerase as described above, examined on a 2% agarose gel to ensure a single band/PCR product and purified with the Promega Wizard SV Gel and PCR Cleanup System (Promega, A9282) Samples are sequenced by Azenta and analyzed using the Poly Peak Parser software available online at http://yosttools.genetics.utah.edu/PolyPeakParser/ ^48^.

### Allele Specific PCR and Restriction Enzyme Analysis of F1 Adults

DNA collected from the tails of F1 fish was PCR amplified with allele specific forward primers and wild-type reverse primers (Table S2) using the Phusion Hot Start II DNA Polymerase and PCR protocols as described above. PCR products were examined on a 2% agarose gel to verify the presence or absence of a band. For restriction enzyme analysis, PCR products were amplified as described above with the Phusion Hot Start II DNA Polymerase. 8ul of each PCR product was then incubated with 2ul of Cut Smart Buffer, 1ul of XbaI, and water up to 20ul at 37°C for one hour. Digested and undigested PCR products were examined on a 2% agarose gel to identify those samples that underwent restriction enzyme digestion.

## ACKNOWLEDGMENTS

The authors would like to acknowledge the members of the Parant lab for technical help and critical reading of the manuscript. Also, NIH ORIP and the UAB Center for Precision Animal Modeling (CPAM) for supporting generation of zebrafish patient alleles. We use the following core facilities: UAB Zebrafish Research Facility and UAB Heflin Center for Genomic Science. JMP, MSA, and BKY are supported by NIH U54OD030167. JMP and BKY are supported by NIH U54 DK126087. JMP is also supported by NIH R01CA216108.

## AUTHOR CONTRIBUTIONS

JMP, MSA, BKY and HRT designed and supervised the study. HRT generated zebrafish mutants. HRT in consultation with JMP, made all figures and performed statistical analyses. HRT and JMP wrote the manuscript.

## DECLARATION OF GENERATIVE AI AND AI-ASSISTED TECHNOLOGIES

AI was not used.

## SUPPLEMENTAL INFORMATION

Figures S1–S5 and Table S1 and S2

Figure S1: Guide design and evaluation.

Figure S2: Indels observed by NGS and ICE analysis.

Figure S3: False positives through ICE analysis.

Figure S4: HRMA detection of adult F1 patient-carrying variants.

Figure S5: Summary of HDR frequency in 12 projects at different stages of production.

Table S1: HDR oligos used.

Table S2: Primers used for PCR

