## Supplementary figures and tables for "Engineering precision zebrafish alleles of human disease"

**Supplemental figure legends:**

**Figure S1: guide design and evaluation.** Guide design and HRMA evaluation for A) *nf1a* GR, B) *vma21* AV, C) *spop* DN, D) *sgo1* KE, E) *pex10* HD, and F) *fkrp* CY projects.

**Figure S2: indels observed by NGS and ICE analysis**. The prominent alleles identified and frequency by NGS and ICE in pooled 5dpf embryos injected with the oligo plus RNP mixture.

**Figure S3: False positives through ICE analysis.** A) ICE false positives in analysis of F0 progeny with *nf1a* GR alteration. B) ICE false positives in analysis of F0 progeny with *fkrp* CY alteration. C) ICE false positives in analysis of F0 progeny with *fkrp* CY alteration but never contributed to F1 adults or by subsequent NGS was negative. D) ICE false positives in *nf1a* R* projects.

**Figure S4: HRMA detection of adult F1 patient-carrying variants**. A) Example of HRM analysis of F1 adult tail biopsies from *nf1a* RQ F0#7 (15.7% HDR rate in F1 embryos). Different curves and associated allelic sequence are denoted by colored curves. Below are Sanger sequencing chromatogram of gDNA from F1 animals with different curves; along with the frequency of the allele in 42 F1 analyzed and the pooled F1 embryos NGS data. B) Example of HRM analysis of F1 adult tail biopsies from *vma21* AV (19.4% HDR rate in F1 embryos). Different curves and associated allelic sequence are denoted by colored curves. Below are Sanger sequencing chromatogram of gDNA from F1 animals with different curves; along with the frequency of the allele in 53 F1 analyzed and the pooled F1 embryos NGS data. C) Example of HRM analysis of F1 adult tail biopsies from *spop* DN (21.3% HDR rate in F1 embryos). Different curves and associated allelic sequence are denotes by colored curves. Below are Sanger sequencing chromatogram of gDNA from F1 animals with different curves; along with the frequency of the allele in 41 F1 analyzed and the pooled F1 embryos NGS data.

**Figure S5: Summary of HDR frequency in 12 projects at different stages of production**.

**Table S1:** HDR oligos used.

**Table S2:** Primers used for PCR

### Supplemental Figure 1

## A

```
                 c.2542G>A: G848R
Hs   NF1: ATGACTGGCTTCCTTTGTGCCCTTGGGGGAGTGTGCCTCCAGCAGAGAAGC
Dr nf1a: ATGACTGGTTTCTTGTGCGCTCTGGGAGGTGTTTGTCTCCAACAGCGCAGC
         -M--T--G--F--L--C--A--L--G--G--V--C--L--Q--Q--R--S-
                 c.2287G>A: G763R
```

Guide 1:

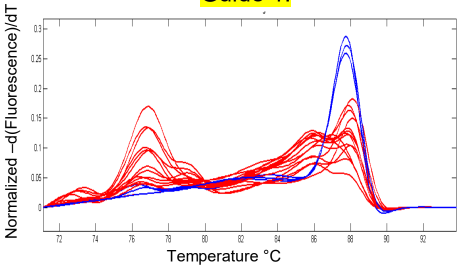


Guide 2:

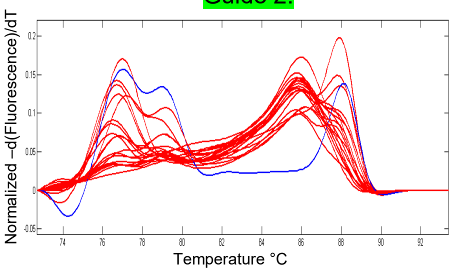


Guide 3:

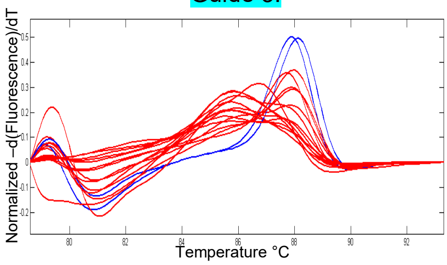


## B

```
                              c.164G>T: G55V
Hs VMA21: aaactgtttttttttctcttgatagGCGCCCTTGGGATGTCCAATAGGGAC
Dr vma21: tcttgcttatttctgtcttcgcagCCACTTTGGGCTACTCCAGCAATGAC
                                  A--T--L--G--Y--S--S--N--D-
                             c.173C>T: A58V
```

Guide 1:

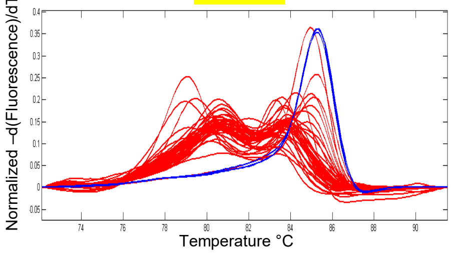


## C

```
                              c.430G>A: D144N
Hs SPOP: TTCATCCGTAGAGATTTTCTTTTGGATGAGGCCAACGGGCTTCTCCCTGAT
Dr spop: TTCATAAGGCGAGACTTTCTTCTAGATGAGGCCAACGGGCTGCTGCCTGAT
         -F--I--R--R--D--F--L--L--D--E--A--N--G--L--L--P--D-
                              c.430G>A: D144N
```

Guide 1:

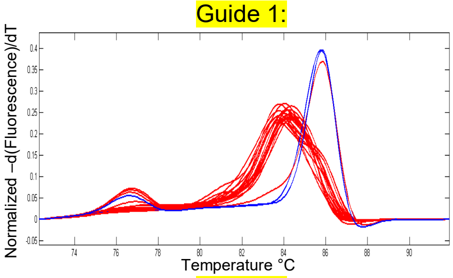


## D

```
                               c.67A>G: K23E
Hs SGO1: CTTGAAGACATAAAGAAGCGAATGAAAGAGAAAAGGAATAAAAACTTGGCA
Dr sgo1: CTGGAGGACAATAAAGAGAAGATGAAGGAGAAGCGGAACCGGCGGCTGAGC
         -L--E--D--N--K--E--K--M--K--E--K--R--N--R--R--L--S-
                               c.55A>G: K19E
```

Guide 1:

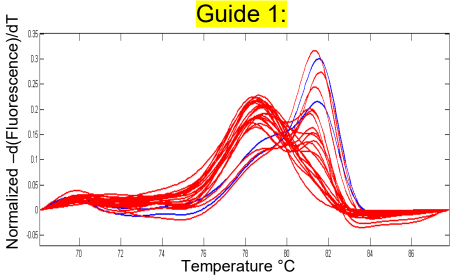


## E

```
                                c.928C>G: H310D
Hs PEX10: CACCCAACAGCCACGCCCTGCGGCCACCTGTTCTGCTGGGAGTGCATCACC
Dr pex10: AACACCACCAGCACTCCCTGCGGCCATCTTTTCTGCTGGGAGTGCATTACC
          -N--T--T--S--T--P--C--G--H--L--F--C--W--E--C--I--T-
                                c.844C>G: H282D
```

Guide 1:

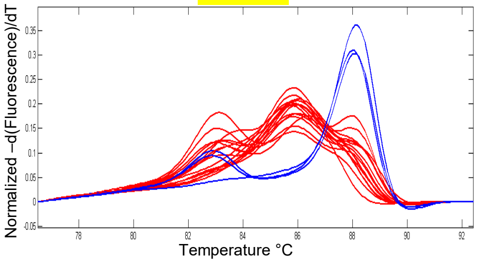


Guide 2:

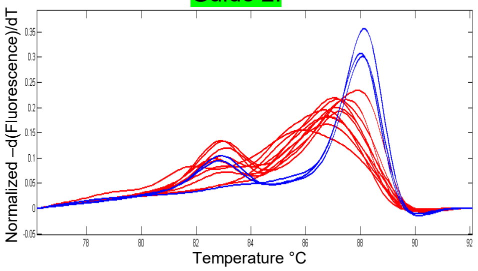

F

**c.953G>A: C318Y**

```
Hs FKRP: GAGGAGCGCTGGACGCCCCCCTGCTGCCTGCGCGCGCTGCGCGAGACCGCC
Dr fkrp: ATTGAGCGCTGGACTCCACCTTGCTGCTTACGTGCCCTCAGAGAGACCACT
         -I--E--R--W--T--P--P--C--C--L--R--A--L--R--E--T--T-
```

**c.1064G>A: C355Y**

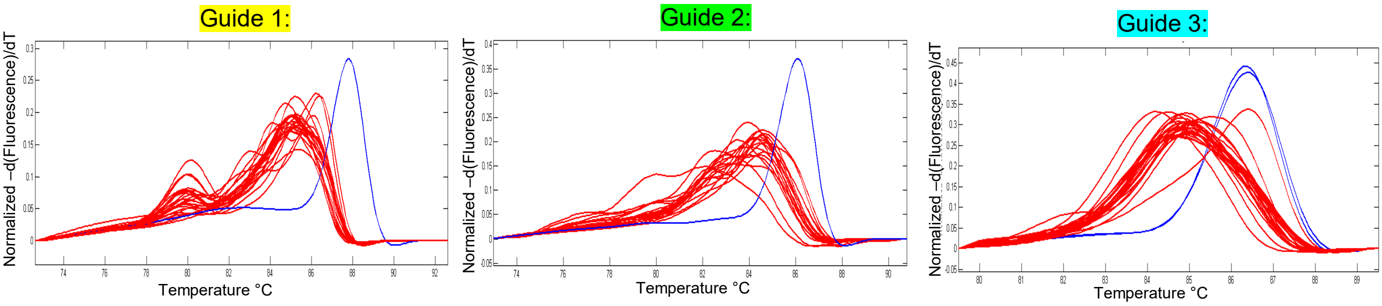

### Supplemental Figure 2

| | **Alleles Detected by NGS** | **Alleles Detected by ICE Analysis** |
|---|---|---|
| *nf1a* R1191Q | Δ2 (-TT): 4.5%<br>R1191R HDR: 4.2%<br>Δ4 (-TCTT): 1.7%<br>Δ1 (-T): 1.5%<br>Δ5 (-TCTTT): 1.2% | Δ2 (-TT): 5%<br>Δ1 (-T): 2%<br>R1191R HDR: 1%<br>Δ4 (-TTTA): 1%<br>Δ5 (-TCTTT): 1% |
| *nf1a* G763R | Δ8 (-CTTGTGCG): 4%<br>Δ5 (-GCTCT): 3.2%<br>Δ2 (-GC): 2.7%<br>G763R HDR: 2.2%<br>Δ7 (-TGCGCTC): 1.8% | Δ4 (-GCTC): 4%<br>Δ2 (-GC): 4%<br>Δ1 (-): 4%<br>G763R HDR: 1%<br>+4 (+NNNN): 1% |
| *vma21* A58V | Δ7 (-CTTTGGG): 9.4%<br>Δ10 (-ACTTTGGGC): 8.5%<br>Δ2 (TT): 4.9%<br>A58V HDR: 4.3%<br>Δ10 (-GCCACTTTGG): 3.8% | A58V HDR: 8%<br>Δ10 (-CTACTCCAGC): 8%<br>Δ7 (-CTCCAGC): 8%<br>Δ2 (-GC): 3%<br>+3 (+NNN): 2% |
| *spop* D144N | Δ8 (-GGCCAACG): 18.7%<br>Δ7 (-GGCCAAC): 7.9%<br>D144N HDR: 3.9%<br>+1 (+C): 3.8%<br>Δ1 (-C): 3.4% | Δ8 (-AGGCCAAC): 16%<br>Δ7 (-GAGGCCA): 10%<br>Δ8 (-GGCCAACG): 6%<br>+1 (+N): 5%<br>Δ2 (-CC): 5%<br>NOTE: D144N HDR: NOT DETECTED |
| *sgo1 K19E* | Δ6 (-GAAGAT): 14.5%<br>Δ12 (-GAGAAGATGAAG): 11.7%<br>Δ3 (-GAT): 9.7%<br>K19E HDR: 3.2%<br>+1 (+A): 2.4% | Δ3 (-ATG): 17%<br>K19E HDR: 14%<br>Δ6 (-ATGAAG): 12%<br>Δ12 (-GAGAAGATGAAG): 8%<br>Δ12 (-ATGAAGGAGAAG): 5% |
| *pex10* H282D | Δ3 (-GCG): 17.6%<br>Δ7 (-CCTGCGG): 4.2%<br>Δ15 (-CTGCGGCCATCTTTT): 1.5%<br>H282D HDR: 1.5%<br>Δ3 (-GCC): 1.3% | Δ3 (-GCG): 16%<br>Δ3 (-CGG): 15%<br>Δ7 (-CGGCCAT): 4%<br>H282D HDR: 2%<br>Δ15 (-CTGCGGCCATCTTTT): 2% |
| *fkrp* C355Y | Δ4 (-CTTG): 11.9%<br>Δ7 (-CTTGCTG): 5.1%<br>Δ1 (-T): 4.3%<br>+1 (+T): 4.2%<br>C355Y HDR: 1.4% | C355Y HDR: 58%<br>Δ4 (-GTGC): 13% |

### Supplemental Figure 3

A) ICE indicated correct HDR, when only positive for 1 of 4 HDR base changes.

```
** nf1a wt:                      TTTCTTGTGCGCTCTGGGAGGTGTTTGTCT
   nf1a GR desired HDR:          TTTCTTGTGCGCCTTAAGAGGTGTTTGTCT
   nf1a GR ICE false positive:   TTTCTTGTGCGCTCTGAGAGGTGTTTGTCT
```

B) ICE indicated correct *fkrp* CY HDR in individual F1 progeny, but did not detect extra SNP nearby HDR mutation.

```
fkrp wt  :                CCACCTTGCTGCTTACGTGCCCTCAGAGAGA
fkrp CY desired HDR:      CCACCTTGCTACTTACGTGCGCTCAGAGAGA
fkrp CY with extra SNP:   TCACCTTGCTACTTACGTGCGCTCAGAGAGA
```

C) ICE indicated HDR positive in pooled *fkrp* CY F1 gDNA, but HDR not present in progeny or NGS data.

1. F0 #3: ICE HDR rate was 6% but NGS indicated 0% HDR
2. F0 #4: ICE HDR rate was 2% but NGS indicated 0% HDR
3. F0 #11: ICE HDR rate was 3% but mutation not present in 121 F1 embryos
   wild-type: 59 of 121 (49%); SNPs: 15 of 121 (12%), Δ16/+: 18 of 121 (15%), +1/+: 24 of 121 (20%), and Δ30/+: 5 of 121 (4%)
4. F0#12: ICE HDR rate was 2% but NGS indicated 0% HDR

D) ICE indicated HDR positive in 6 of 6 *nf1* R* F1 pooled progeny gDNA; further analyzed 3 of the 6 ICE positive F0s and only 1 of the 3 produced progeny that have the HDR allele by Sanger sequencing.

1. F0 #1 (good F0):
   ICE analysis of F1 pooled gDNA: HDR (3%), +1/+ (25%), +9/+ (1%)
   Sanger sequencing of individual F1: HDR (5%: 3/62), +1/+: (24%: 15/62), Δ10/+ (18%:11/62), +9/+ (16%: 10/62)
2. F0 #2 (false positive by ICE):
   ICE analysis of F1 pooled gDNA: HDR (14%), Δ6/+ (1%)
   Sanger sequencing of individual F1: +1/+ (1.4%: 1/72 ), +24/+ (40%: 29/72), Δ6/+ (11%: 8/72), and 0% HDR
3. F0 #3 (false positive by ICE):
   ICE analysis of F1 pooled gDNA: HDR (4%), Δ10/+[1] (27%), +1/+ (12%), Δ10[2] (10%), Δ10[3] (3%)
   Sanger sequencing of individual F1: +1/+ (42%: 34/81), Δ10/+ (47%: 38/81), and 0% HDR

### Supplemental Figure 4

A

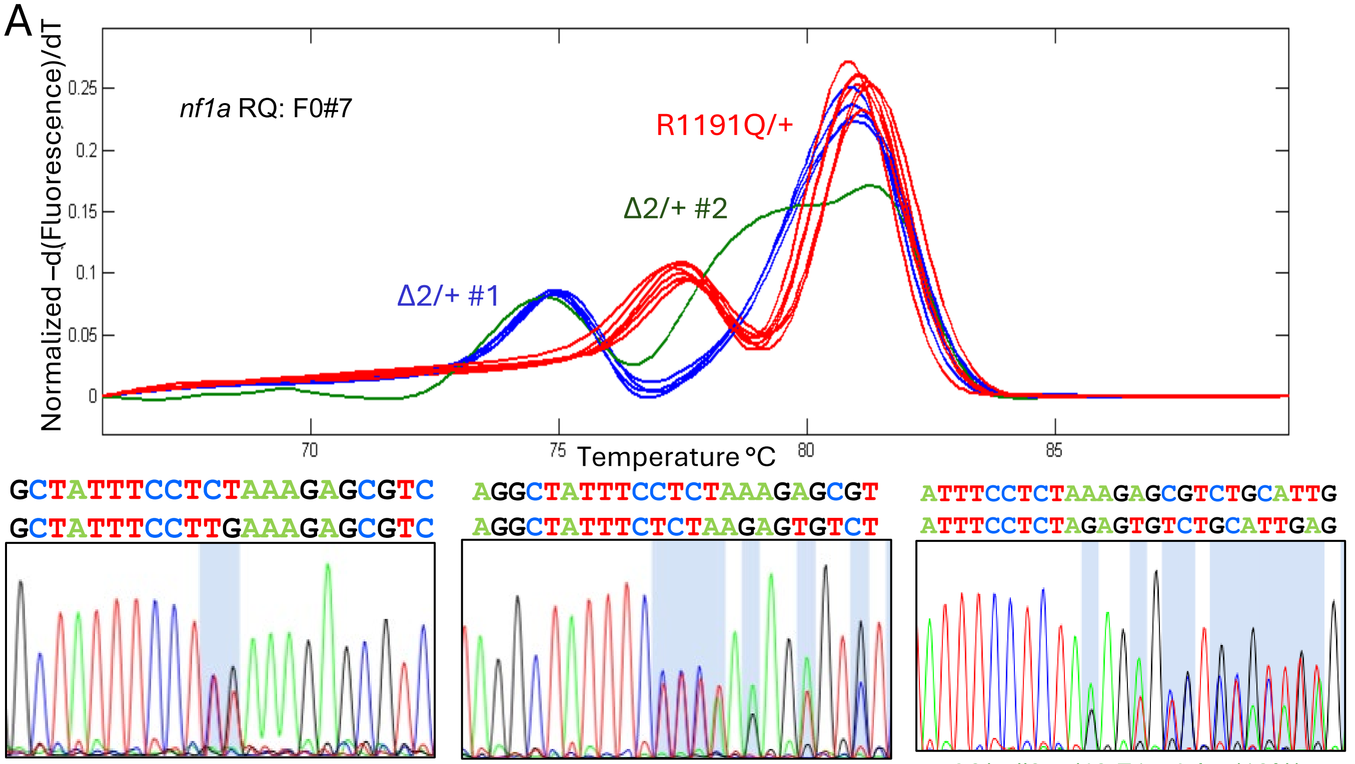


RQ/+: 8/42 F1 adults (19%) - NGS frequency was 15.7%

Δ2/+ #1: 9/42 F1 adults (21%) – NGS frequency was 16%

Δ2/+ #2: 5/42 F1 adults (12%) – NGS frequency was 6.8%

B

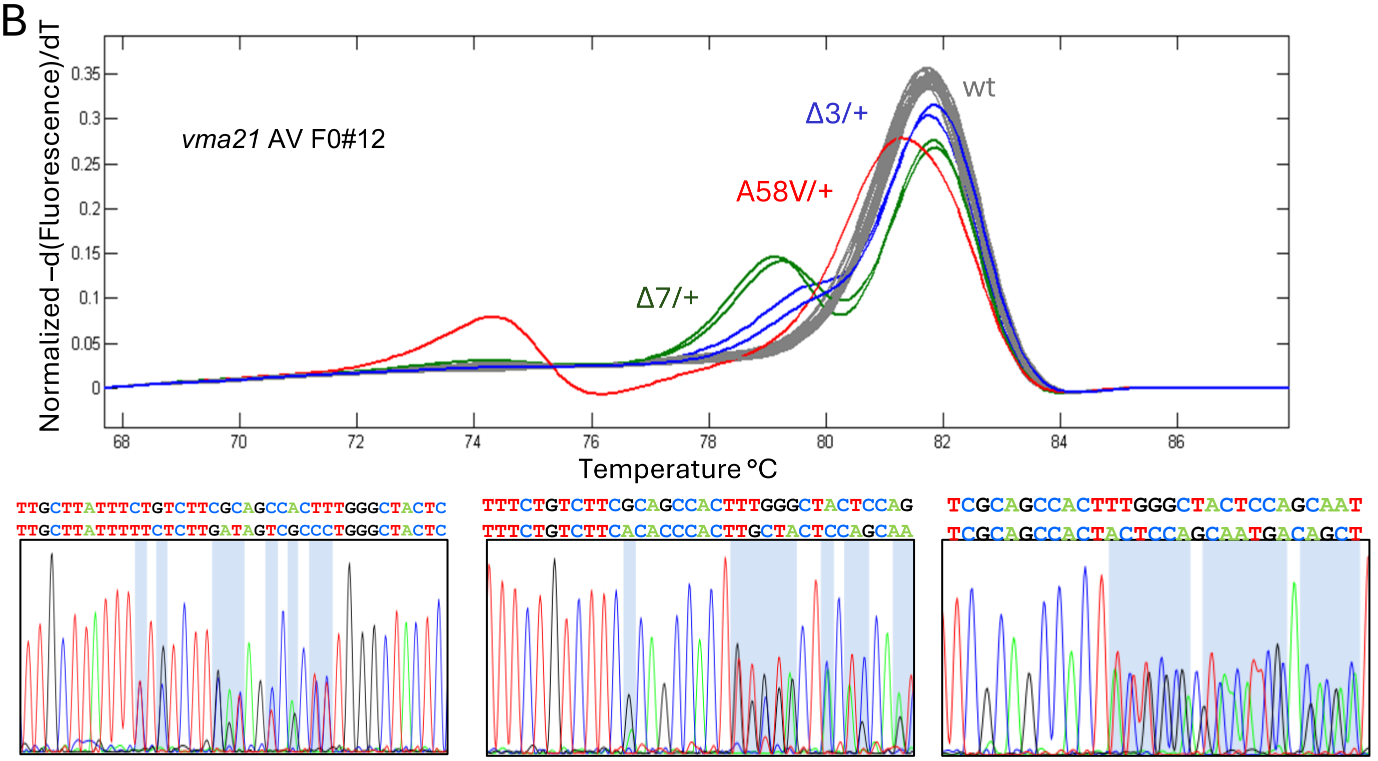


AV/+: 6/53 F1 adults (11%) – NGS frequency was 19.4%

Δ3/+: 6/53 F1 adults (11%) – NGS frequency was 0% (not detected)

Δ7/+: 1/53 F1 adults (2%) – NGS frequency was 0% (not detected)

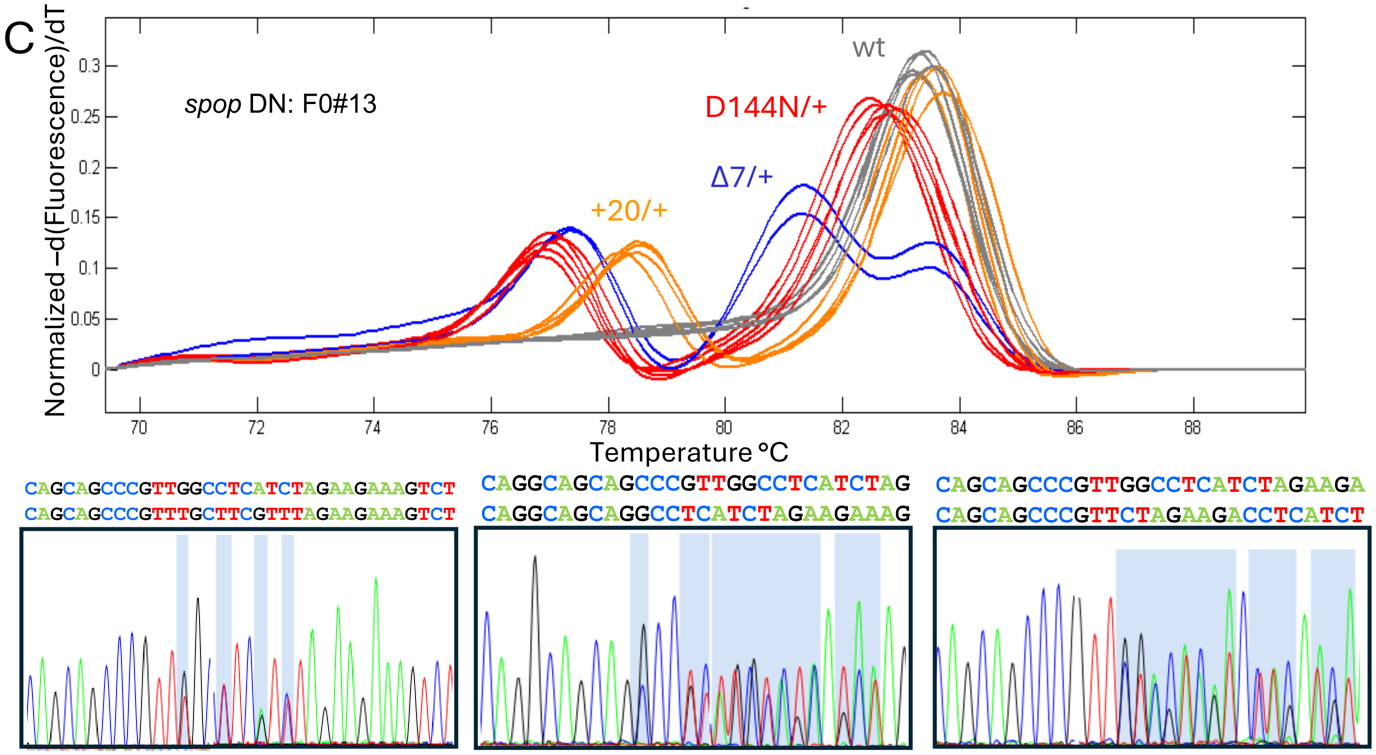


D144N/+: 8/41 F1 adults(19.5%) - NGS frequency was 21.3%

Δ7/+: 4/41 F1 adults (9.8%) - NGS frequency was 1%

+20/+: 7/41 F1 adults (17%) – NGS frequency was 7.5%

### Supplemental Figure 5

| Allele | Injected F0 Embryo NGS HDR Frequency | F0 + Frequency (>2% GTF) | NGS HDR Frequency in F1 pooled embryos from positive F0 | Mutation Frequency in F1 Adults |
|---|---|---|---|---|
| *nf1a R1191Q* | 4.2 | 1 of 12 (8.3%) | 15.7% | 8 of 42 (19%) |
| *nf1a G763R* | 2.2 | 3 of 35 (8.6%) | 3.3, 4.1, 22.7% | 6 of 38 (15.8%) |
| *vma21 A58V* | 4.3 | 3 of 16 (18.8%) | 5.2, 14, 19.4% | 6 of 53 (11.3%) |
| *spop D144N* | 3.9 | 2 of 16 (12.5%) | 6.6, 21.3% | 8 of 41 (19.5%) |
| sgo1 *K19E* | 3.2 | 3 of 11 (27.3%) | 2.7, 4.5, 18.3% | >4 of 60 (>6.7%) |
| *pex10 H282D* | 1.5 | 1 of 23 (4.4%) | 6.1% | 5 of 96 (5.2%) |
| *fkrp C355Y* | 2.6 with G3<br>1.4 with G1 | 0 of 40 (0%)<br>1 of 17 (5.9%) | 2.1% | 6 of 94 (6.4%) |
| *nf1a* R628* | 6.1 | NA | NA | 4 of 54 (7.4%) |
| *nf1a* M907del | 2.8 | 1 of 4 (25%) | 17.1% | 9 of 48 (18.8%) |
| *tp53* R144H | 10.6 | 2 of 14 (14.3%) | 18.6, 6.8% | 8 of 57 (14%) |
| *pkd2* L593W | 3.8 | 2 of 12 (16.7%) | 6.3, 2.5% | 2 of 39 (5.1%) |
| *tp53* K89R | 3 | 3 of 12 (25%) | 5.2, 5.6, 3% | 5 of 72 (6.9%) |

### Supplemental Table 1

| ***nf1a* R1191Q with Guide 1 – 0 changes** | |
|---|---|
| Comp-asymmetrical 90:36 | TCTCTTCGACTCTCGCCATCTGCTGTATCAGCTGTTGTGGAATATGTTCTCCAAAGAGGTGGAGCTGGCAGACTCAATGCAGACGCTCTTTCAAGGAAATAGCCTAGCCAGCAAGATTATGACCTT |
| Rev Comp-asymmetrical 90:36 | AAGGTCATAATCTTGCTGGCTAGGCTATTTCCTTGAAAGAGCGTCTGCATTGAGTCTGCCAGCTCCACCTCTTTGGAGAACATATTCCACAACAGCTGATACAGCAGATGGCGAGAGTCGAAGAGA |
| Comp-symmetrical 60:60 | GCTGTTGTGGAATATGTTCTCCAAAGAGGTGGAGCTGGCAGACTCAATGCAGACGCTCTTTCAAGGAAATAGCCTAGCCAGCAAGATTATGACCTTCTGCTTCAAGgtgacgtgaagcgg |
| Rev Comp-symmetrical 60:60 | CcgcttcacgtcacCTTGAAGCAGAAGGTCATAATCTTGCTGGCTAGGCTATTTCCTTGAAAGAGCGTCTGCATTGAGTCTGCCAGCTCCACCTCTTTGGAGAACATATTCCACAACAGC |
| Comp-symmetrical 36:36 | AGAGGTGGAGCTGGCAGACTCAATGCAGACGCTCTTTCAAGGAAATAGCCTAGCCAGCAAGATTATGACCTT |
| Rev Comp-symmetrical 36:36 | AAGGTCATAATCTTGCTGGCTAGGCTATTTCCTTGAAAGAGCGTCTGCATTGAGTCTGCCAGCTCCACCTCT |
| Comp-asymmetrical 36:90 | AGAGGTGGAGCTGGCAGACTCAATGCAGACGCTCTTTCAAGGAAATAGCCTAGCCAGCAAGATTATGACCTTCTGCTTCAAGgtgacgtgaagcggtgacagtcattccatgagttcagccttcattgtttt |
| Rev Comp-asymmetrical 36:90 | AaaacaatgaaggctgaactcatggaatgactgtcaccgcttcacgtcacCTTGAAGCAGAAGGTCATAATCTTGCTGGCTAGGCTATTTCCTTGAAAGAGCGTCTGCATTGAGTCTGCCAGCTCCACCTCT |
| ***nf1a* R1191Q with Guide 1 -6 changes** | |
| Comp-asymmetrical 90:36 | TCTCTTCGACTCTCGCCATCTGCTGTATCAGCTGTTGTGGAATATGTTCTCCAAAGAGGTGGAGCTGGCAGACTCAATGCAGACGCTCTTTCAAGGCAACAGCTTGGCAAGTAAGATTATGACCTT |
| Rev Comp-asymmetrical 90:36 | AAGGTCATAATCTTACTTGCCAAGCTGTTGCCTTGAAAGAGCGTCTGCATTGAGTCTGCCAGCTCCACCTCTTTGGAGAACATATTCCACAACAGCTGATACAGCAGATGGCGAGAGTCGAAGAGA |
| Comp-symmetrical 60:60 | GCTGTTGTGGAATATGTTCTCCAAAGAGGTGGAGCTGGCAGACTCAATGCAGACGCTCTTTCAAGGCAACAGCTTGGCAAGTAAGATTATGACCTTCTGCTTCAAGgtgacgtgaagcgg |
| Rev Comp-symmetrical 60:60 | ccgcttcacgtcacCTTGAAGCAGAAGGTCATAATCTTACTTGCCAAGCTGTTGCCTTGAAAGAGCGTCTGCATTGAGTCTGCCAGCTCCACCTCTTTGGAGAACATATTCCACAACAGC |
| Comp-symmetrical 36:36 | AGAGGTGGAGCTGGCAGACTCAATGCAGACGCTCTTTCAAGGCAACAGCTTGGCAAGTAAGATTATGACCTT |
| Rev Comp-symmetrical 36:36 | AAGGTCATAATCTTACTTGCCAAGCTGTTGCCTTGAAAGAGCGTCTGCATTGAGTCTGCCAGCTCCACCTCT |
| Comp-asymmetrical 36:90 | AGAGGTGGAGCTGGCAGACTCAATGCAGACGCTCTTTCAAGGCAACAGCTTGGCAAGTAAGATTATGACCTTCTGCTTCAAGgtgacgtgaagcggtgacagtcattccatgagttcagccttcattgtttt |
| Rev Comp-asymmetrical 36:90 | aaaacaatgaaggctgaactcatggaatgactgtcaccgcttcacgtcacCTTGAAGCAGAAGGTCATAATCTTACTTGCCAAGCTGTTGCCTTGAAAGAGCGTCTGCATTGAGTCTGCCAGCTCCACCTCT |

| ***nf1a* R1191Q with Guide 3 – 1 change** | |
|---|---|
| Comp-asymmetrical 90:36 | cgcttcacgtcacCTTGAAGCAGAAGGTCATAATCTTGCTTGCTAGGCTATTTCCTTGAAAGAGCGTCTGCATTGAGTCTGCCAGCTCCACCTCTTTGGAGAACATATTCCACAACAGCTGATACA |
| Rev Comp-asymmetrical 90:36 | TGTATCAGCTGTTGTGGAATATGTTCTCCAAAGAGGTGGAGCTGGCAGACTCAATGCAGACGCTCTTTCAAGGAAATAGCCTAGCAAGCAAGATTATGACCTTCTGCTTCAAGgtgacgtgaagcg |
| Comp-symmetrical 60:60 | ctgaactcatggaatgactgtcaccgcttcacgtcacCTTGAAGCAGAAGGTCATAATCTTGCTTGCTAGGCTATTTCCTTGAAAGAGCGTCTGCATTGAGTCTGCCAGCTCCACCTCTTTGGA |
| Rev Comp-symmetrical 60:60 | TCCAAAGAGGTGGAGCTGGCAGACTCAATGCAGACGCTCTTTCAAGGAAATAGCCTAGCAAGCAAGATTATGACCTTCTGCTTCAAGgtgacgtgaagcggtgacagtcattccatgagttcag |
| Comp-symmetrical 36:36 | cgcttcacgtcacCTTGAAGCAGAAGGTCATAATCTTGCTTGCTAGGCTATTTCCTTGAAAGAGCGTCTGCATTGA |
| Rev Comp-symmetrical 36:36 | TCAATGCAGACGCTCTTTCAAGGAAATAGCCTAGCAAGCAAGATTATGACCTTCTGCTTCAAGgtgacgtgaagc |
| Comp-asymmetrical 36:90 | agtgcggtaactcatgcaaaacaatgaaggctgaactcatggaatgactgtcaccgcttcacgtcacCTTGAAGCAGAAGGTCATAATCTTGCTTGCTAGGCTATTTCCTTGAAAGAGCGTCTGCA |
| Rev Comp-asymmetrical 36:90 | TGCAGACGCTCTTTCAAGGAAATAGCCTAGCAAGCAAGATTATGACCTTCTGCTTCAAGgtgacgtgaagcggtgacagtcattccatgagttcagccttcattgttttgcatgagttaccgcact |
| ***nf1a* R1191Q with Guide 3 – 6 changes** | |
| Comp-asymmetrical 90:36 | cgcttcacgtcacCTTGAAGCAGAAGGTCATAATCTTACTTGCCAAGCTGTTGCCTTGAAAGAGCGTCTGCATTGAGTCTGCCAGCTCCACCTCTTTGGAGAACATATTCCACAACAGCTGATACA |
| Rev Comp-asymmetrical 90:36 | TGTATCAGCTGTTGTGGAATATGTTCTCCAAAGAGGTGGAGCTGGCAGACTCAATGCAGACGCTCTTTCAAGGCAACAGCTTGGCAAGTAAGATTATGACCTTCTGCTTCAAGgtgacgtgaagcg |
| Comp-symmetrical 60:60 | ctgaactcatggaatgactgtcaccgcttcacgtcacCTTGAAGCAGAAGGTCATAATCTTACTTGCCAAGCTGTTGCCTTGAAAGAGCGTCTGCATTGAGTCTGCCAGCTCCACCTCTTTGGA |
| Rev Comp-symmetrical 60:60 | TCCAAAGAGGTGGAGCTGGCAGACTCAATGCAGACGCTCTTTCAAGGCAACAGCTTGGCAAGTAAGATTATGACCTTCTGCTTCAAGgtgacgtgaagcggtgacagtcattccatgagttcag |
| Comp-symmetrical 36:36 | cgcttcacgtcacCTTGAAGCAGAAGGTCATAATCTTACTTGCCAAGCTGTTGCCTTGAAAGAGCGTCTGCATTGA |
| Rev Comp-symmetrical 36:36 | TCAATGCAGACGCTCTTTCAAGGCAACAGCTTGGCAAGTAAGATTATGACCTTCTGCTTCAAGgtgacgtgaagcg |
| Comp-asymmetrical 36:90 | agtgcggtaactcatgcaaaacaatgaaggctgaactcatggaatgactgtcaccgcttcacgtcacCTTGAAGCAGAAGGTCATAATCTTACTTGCCAAGCTGTTGCCTTGAAAGAGCGTCTGCA |
| Rev Comp-asymmetrical 36:90 | TGCAGACGCTCTTTCAAGGCAACAGCTTGGCAAGTAAGATTATGACCTTCTGCTTCAAGgtgacgtgaagcggtgacagtcattccatgagttcagccttcattgttttgcatgagttaccgcact |

| ***nf1a* G763R with Guide 1 -0 changes** | |
|---|---|
| Comp-asymmetrical 90:36 | tggtcaattatttcatcttttttccattcatggtcatctattttctgttcgtcctcagGAGTGGATCAACATGACTGGTTTCTTGTGCGCTCTGAGAGGTGTTTGTCTCCAACAGCGCAGCACGC |
| Rev Comp-asymmetrical 90:36 | GCGTGCTGCGCTGTTGGAGACAAACACCTCTCAGAGGCGCACAAGAAACCAGTCATGTTGATCCACTCctgaggacgaacagaaaatagatgaccatgaatggaaaaaaagatgaaataattgacca |
| Comp-symmetrical 60:60 | atggtcatctattttctgttcgtcctcagGAGTGGATCAACATGACTGGTTTCTTGTGCGCTCTGAGAGGTGTTTGTCTCCAACAGCGCAGCACGCCGAGTCTGGGCACCTACAGTCCTC |
| Rev Comp-symmetrical 60:60 | GAGGACTGTAGGTGCCCAGACTCGGCGTGCTGCGCTGTTGGAGACAAACACCTCTCAGAGGCGCACAAGAAACCAGTCATGTTGATCCACTCctgaggacgaacagaaaatagatgaccat |
| Comp-symmetrical 36:36 | ctcagGAGTGGATCAACATGACTGGTTTCTTGTGCGCTCTGAGAGGTGTTTGTCTCCAACAGCGCAGCACGC |
| Rev Comp-symmetrical 36:36 | GCGTGCTGCGCTGTTGGAGACAAACACCTCTCAGAGGCGCACAAGAAACCAGTCATGTTGATCCACTCctgag |
| Comp-asymmetrical 36:90 | ctcagGAGTGGATCAACATGACTGGTTTCTTGTGCGCTCTGAGAGGTGTTTGTCTCCAACAGCGCAGCACGCCGAGTCTGGGCACCTACAGTCCTCCAATGGGCCCCCTGACCGAACGCAAGGGGT |
| Rev Comp-asymmetrical 36:90 | ACCCCTTGCGTTCGGTCAGGGGGCCCATTGGAGGACTGTAGGTGCCCAGACTCGGCGTGCTGCGCTGTTGGAGACAAACACCTCTCAGAGGCGCACAAGAAACCAGTCATGTTGATCCACTCctgag |
| ***nf1a* G763R with Guide 1 -3 changes** | |
| Comp-asymmetrical 90:36 | tggtcaattatttcatcttttttccattcatggtcatctattttctgttcgtcctcagGAGTGGATCAACATGACTGGTTTCTTGTGCGCCTTAAGAGGTGTTTGTCTCCAACAGCGCAGCACGC |
| Rev Comp-asymmetrical 90:36 | GCGTGCTGCGCTGTTGGAGACAAACACCTCTTAAGGCGCACAAGAAACCAGTCATGTTGATCCACTCctgaggacgaacagaaaatagatgaccatgaatggaaaaaaagatgaaataattgacca |
| Comp-symmetrical 60:60 | AtggtcatctattttctgttcgtcctcagGAGTGGATCAACATGACTGGTTTCTTGTGCGCCTTAAGAGGTGTTTGTCTCCAACAGCGCAGCACGCCGAGTCTGGGCACCTACAGTCCTC |
| Rev Comp-symmetrical 60:60 | GAGGACTGTAGGTGCCCAGACTCGGCGTGCTGCGCTGTTGGAGACAAACACCTCTTAAGGCGCACAAGAAACCAGTCATGTTGATCCACTCctgaggacgaacagaaaatagatgaccat |
| Comp-symmetrical 36:36 | ctcagGAGTGGATCAACATGACTGGTTTCTTGTGCGCCTTAAGAGGTGTTTGTCTCCAACAGCGCAGCACGC |
| Rev Comp-symmetrical 36:36 | GCGTGCTGCGCTGTTGGAGACAAACACCTCTTAAGGCGCACAAGAAACCAGTCATGTTGATCCACTCctgag |
| Comp-asymmetrical 36:90 | ctcagGAGTGGATCAACATGACTGGTTTCTTGTGCGCCTTAAGAGGTGTTTGTCTCCAACAGCGCAGCACGCCGAGTCTGGGCACCTACAGTCCTCCAATGGGCCCCCTGACCGAACGCAAGGGGT |
| Rev Comp-asymmetrical 36:90 | ACCCCTTGCGTTCGGTCAGGGGGCCCATTGGAGGACTGTAGGTGCCCAGACTCGGCGTGCTGCGCTGTTGGAGACAAACACCTCTTAAGGCGCACAAGAAACCAGTCATGTTGATCCACTCctgag |

| *vma21* A58V with Guide 1 – 3 changes | |
|---|---|
| Comp-asymmetrical 90:36 | CAGCGTAGAAGTAGCTGTCATTGCTGGAGTAGCCCAAAGCGActgcgaagacagaaataagcaagaatgatcaatgattactaaagcaaatcaaaaacactgcacctaaagacaggagtactctgt |
| Rev Comp-asymmetrical 90:36 | acagagtactcctgtctttaggtgcagtgtttttgatttgctttagtaatcattgatcattcttgcttatttctgtcttcgcagTCGCTTTGGGCTACTCCAGCAATGACAGCTACTTCTACGCTG |
| ***vma21* A58V with Guide 1 – 8 changes** | |
| Comp-asymmetrical 90:36 | CAGCGTAGAAGTAGCTGTCATTGCTGGAGTAGCCCAGGGCGActatcaagagaaaaataagcaagaatgatcaatgattactaaagcaaatcaaaaacactgcacctaaagacaggagtactctgt |
| Rev Comp-asymmetrical 90:36 | acagagtactcctgtctttaggtgcagtgtttttgatttgctttagtaatcattgatcattcttgcttattttctcttgatagTCGCCCTGGGCTACTCCAGCAATGACAGCTACTTCTACGCTG |
| Comp-symmetrical 60:60 | GGACGGCAAGCACAGCGAGGATGGCAGCGTAGAAGTAGCTGTCATTGCTGGAGTAGCCCAGGGCGActatcaagagaaaaataagcaagaatgatcaatgattactaaagcaaatcaaaa |
| Rev Comp-symmetrical 60:60 | ttttgatttgctttagtaatcattgatcattcttgcttattttctcttgatagTCGCCCTGGGCTACTCCAGCAATGACAGCTACTTCTACGCTG CCATCCTCGCTGTGCTTGCCGTCC |
| Comp-symmetrical 36:36 | CCAGCGTAGAAGTAGCTGTCATTGCTGGAGTAGCCCAGGGCGActatcaagagaaaaataagcaagaatgatc |
| Rev Comp-symmetrical 36:36 | gatcattcttgcttattttctcttgatagTCGCCCTGGGCTACTCCAGCAATGACAGCTACTTCTACGCTG |
| Comp-asymmetrical 36:90 | CCACATAAACGAAGAGCGCCAGAACCACATGGACGGCAAGCACAGCGAGGATGGCAGCGTAGAAGTAGCTGTCATTGCTGGAGTAGCCCAGGGCGActatcaagagaaaataagcaagaatgatc |
| Rev Comp-asymmetrical 36:90 | gatcattcttgcttattttctcttgatagTCGCCCTGGGCTACTCCAGCAATGACAGCTACTTCTACGCTGCCATCCTCGCTGTGCTTGCCGTCCATGTGGTTCTGGCGCTCTTCGTTTATGTGG |

| ***spop* D144N with Guide 1 – 3 changes** | |
|---|---|
| Comp-asymmetrical 90:36 | gcagAGAGCCAACGGGCATATCGCTTCGTGCAAGGGAAAGACTGGGGTTTCAAGAAGTTCATAAGGCGAGACTTTCTTCTAAACGATGCAAACGGGCTGCTGCCTGATGACAAGTTAACGCTCTTC |
| Rev Comp-asymmetrical 90:36 | GAAGAGCGTTAACTTGTCATCAGGCAGCAGCCCGTTTGCATCGTTTAGAAGAAAGTCTCGCCTTATGAACTTCTTGAAACCCCAGTCTTTCCCTTGCACGAAGCGATATGCCCGTTGGCTCTctgc |
| Comp-symmetrical 60:60 | CAAGGGAAAGACTGGGGTTTCAAGAAGTTCATAAGGCGAGACTTTCTTCTAAACGATGCAAACGGGCTGCTGCCTGATGACAAGTTAACGCTCTTCTGTGAGgtaggtgatgtggtcaat |
| Rev Comp-symmetrical 60:60 | attgaccacatcacctacCTCACAGAAGAGCGTTAACTTGTCATCAGGCAGCAGCCCGTTTGCATCGTTTAGAAGAAAGTCTCGCCTTATGAACTTCTTGAAACCCCAGTCTTTCCCTTG |
| Comp-symmetrical 36:36 | AAGTTCATAAGGCGAGACTTTCTTCTAAACGATGCAAACGGGCTGCTGCCTGATGACAAGTTAACGCTCTTC |
| Rev Comp-symmetrical 36:36 | GAAGAGCGTTAACTTGTCATCAGGCAGCAGCCCGTTTGCATCGTTTAGAAGAAAGTCTCGCCTTATGAACTT |
| Comp-asymmetrical 36:90 | AAGTTCATAAGGCGAGACTTTCTTCTAAACGATGCAAACGGGCTGCTGCCTGATGACAAGTTAACGCTCTTCTGTGAGgtaggtgatgtggtcaatcatttactgtgctttccatcaccggtttaa |
| Rev Comp-asymmetrical 36:90 | ttaaaccggtgatggaaagcacagtaaatgattgaccacatcacctacCTCACAGAAGAGCGTTAACTTGTCATCAGGCAGCAGCCCGTTTGCATCGTTTAGAAGAAAGTCTCGCCTTATGAACTT |

| ***sgo1* K19E with Guide 1 – 1 changes** | |
|---|---|
| Comp-asymmetrical 90:36 | ACTGCGGGTGTTTGAGGCGGTGGTGACCGGAGGCAACGATGGTGCAGAAGAAGAGCTTCCAGCAGAGTCTGGAGGACAATAAAGAGAAGATGGAAGAGAAGCGGAACCGGCGGCTGAGCGCGGCGG |
| Rev Comp-asymmetrical 90:36 | CCGCCGCGCTCAGCCGCCGGTTCCGCTTCTCTTCCATCTTCTCTTTATTGTCCTCCAGACTCTGCTGGAAGCTCTTCTTCTGCACCATCGTTGCCTCCGGTCACCACCGCCTCAAACACCCGCAGT |
| Comp-symmetrical 60:60 | AGGCAACGATGGTGCAGAAGAAGAGCTTCCAGCAGAGTCTGGAGGACAATAAAGAGAAGATGGAAGAGAAGCGGAACCGGCGGCTGAGCGCGGCGGCTTCGGCCCGGAGATCAGTGCAGC |
| Rev Comp-symmetrical 60:60 | GCTGCACTGATCTCCGGGCCGAAGCCGCCGCGCTCAGCCGCCGGTTCCGCTTCTCTTCCATCTTCTCTTTATTGTCCTCCAGACTCTGCTGGAAGCTCTTCTTCTGCACCATCGTTGCCT |
| Comp-symmetrical 36:36 | GCTTCCAGCAGAGTCTGGAGGACAATAAAGAGAAGATGGAAGAGAAGCGGAACCGGCGGCTGAGCGCGGCGG |
| Rev Comp-symmetrical 36:36 | CCGCCGCGCTCAGCCGCCGGTTCCGCTTCTCTTCCATCTTCTCTTTATTGTCCTCCAGACTCTGCTGGAAGC |
| Comp-asymmetrical 36:90 | GCTTCCAGCAGAGTCTGGAGGACAATAAAGAGAAGATGGAAGAGAAGCGGAACCGGCGGCTGAGCGCGGCGGCTTCGGCCCGGAGATCAGTGCAGCTCCGGGGAGCAGgtcggtggagagacgctc |
| Rev Comp-asymmetrical 36:90 | gagcgtctctccaccgacCTGCTCCCCGGAGCTGCACTGATCTCCGGGCCGAAGCCGCCGCGCTCAGCCGCCGGTTCCGCTTCTCTTCCATCTTCTCTTTATTGTCCTCCAGACTCTGCTGGAAGC |

| | |
|---|---|
| ***pex10* H282D with Guide 1 – 2 changes** | |
| Comp-asymmetrical 90:36 | ACCACTCGGTAATGCACTCCCAGCAGAACAGGTCGCCGCAGGGAGTGCTGGTGGTGTTTCTGCGCTCCTCCAAACACAAGATACAGCGTGAAGTGCGAGACGACGACTGACTGACCTGATGACTGG |
| Rev Comp-asymmetrical 90:36 | CCAGTCATCAGGTCAGTCAGTCGTCGTCTCGCACTTCACGCTGTATCTTGTGTTTGGAGGAGCGCAGAAACACCACCAGCACTCCCTGCGGCGACCTGTTCTGCTGGGAGTGCATTACCGAGTGGT |
| Comp-symmetrical 60:60 | atgaaggtcttgcCTTGGTATTGCACCACTCGGTAATGCACTCCCAGCAGAACAGGTCGCCGCAGGGAGTGCTGGTGGTGTTTCTGCGCTCCTCCAAACACAAGATACAGCGTGAAGTGC |
| Rev Comp-symmetrical 60:60 | GCACTTCACGCTGTATCTTGTGTTTGGAGGAGCGCAGAAACACCACCAGCACTCCCTGCGGCGACCTGTTCTGCTGGGAGTGCATTACCGAGTGGTGCAATACCAAGgcaagaccttcat |
| Comp-symmetrical 36:36 | ACCACTCGGTAATGCACTCCCAGCAGAACAGGTCGCCGCAGGGAGTGCTGGTGGTGTTTCTGCGCTCCTCCA |
| Rev Comp-symmetrical 36:36 | TGGAGGAGCGCAGAAACACCACCAGCACTCCCTGCGGCGACCTGTTCTGCTGGGAGTGCATTACCGAGTGGT |
| Comp-asymmetrical 36:90 | ttaaacttcacagcatcacttcacagctatatgaaggtcttgcCTTGGTATTGCACCACTCGGTAATGCACTCCCAGCAGAACAGGTCGCCGCAGGGAGTGCTGGTGGTGTTTCTGCGCTCCTCCA |
| Rev Comp-asymmetrical 36:90 | TGGAGGAGCGCAGAAACACCACCAGCACTCCCTGCGGCGACCTGTTCTGCTGGGAGTGCATTACCGAGTGGTGCAATACCAAGgcaagaccttcatatagctgtgaagtgatgctgtgaagtttaa |
| ***pex10* H282D with Guide 2 – 2 changes** | |
| Comp-asymmetrical 90:36 | ACCACTCGGTAATGCACTCCCAGCAGAACAGGTCGCCGCAGGGAGTGCTGGTGGTGTTTCTGCGCTCCTCCAAACACAAGATACAGCGTGAAGTGCGAGACGACGACTGACTGACCTGATGACTGG |
| Rev Comp-asymmetrical 90:36 | CCAGTCATCAGGTCAGTCAGTCGTCGTCTCGCACTTCACGCTGTATCTTGTGTTTGGAGGAGCGCAGAAACACCACCAGCACTCCCTGCGGCGACCTGTTCTGCTGGGAGTGCATTACCGAGTGGT |
| Comp-symmetrical 60:60 | atgaaggtcttgcCTTGGTATTGCACCACTCGGTAATGCACTCCCAGCAGAACAGGTCGCCGCAGGGAGTGCTGGTGGTGTTTCTGCGCTCCTCCAAACACAAGATACAGCGTGAAGTGC |
| Rev Comp-symmetrical 60:60 | GCACTTCACGCTGTATCTTGTGTTTGGAGGAGCGCAGAAACACCACCAGCACTCCCTGCGGCGACCTGTTCTGCTGGGAGTGCATTACCGAGTGGTGCAATACCAAGgcaagaccttcat |
| Comp-symmetrical 36:36 | ACCACTCGGTAATGCACTCCCAGCAGAACAGGTCGCCGCAGGGAGTGCTGGTGGTGTTTCTGCGCTCCTCCA |
| Rev Comp-symmetrical 36:36 | TGGAGGAGCGCAGAAACACCACCAGCACTCCCTGCGGCGACCTGTTCTGCTGGGAGTGCATTACCGAGTGGT |
| Comp-asymmetrical 36:90 | ttaaacttcacagcatcacttcacagctatatgaaggtcttgcCTTGGTATTGCACCACTCGGTAATGCACTCCCAGCAGAACAGGTCGCCGCAGGGAGTGCTGGTGGTGTTTCTGCGCTCCTCCA |
| Rev Comp-asymmetrical 36:90 | TGGAGGAGCGCAGAAACACCACCAGCACTCCCTGCGGCGACCTGTTCTGCTGGGAGTGCATTACCGAGTGGTGCAATACCAAGgcaagaccttcatatagctgtgaagtgatgctgtgaagtttaa |

| | |
|---|---|
| ***fkrp* C355Y with Guide 3 – 1 change** | |
| Comp-asymmetrical 90:36 | TTTCGAGAATGTTGATCACGTATTTAGTGGTCTCTCTGAGCGCACGTAAGTAGCAAGGTGGAGTCCAGCGCTCAATGTAAAGATATTCCGGTGTGTCATCTCGAACTGTCCCGAAACATCGCGCAG |
| Rev Comp-asymmetrical 90:36 | CTGCGCGATGTTTCGGGACAGTTCGAGATGACACACCGGAATATCTTTACATTGAGCGCTGGACTCCACCTTGCTACTTACGTGCGCTCAGAGAGACCACTAAATACGTGATCAACATTCTCGAAA |
| Comp-symmetrical 60:60 | CCAGCCAGTAGCGCACACCAGAGCTTTCGAGAATGTTGATCACGTATTTAGTGGTCTCTCTGAGCGCACGTAAGTAGCAAGGTGGAGTCCAGCGCTCAATGTAAAGATATTCCGGTGTGT |
| Rev Comp-symmetrical 60:60 | ACACACCGGAATATCTTTACATTGAGCGCTGGACTCCACCTTGCTACTTACGTGCGCTCAGAGAGACCACTAAATACGTGATCAACATTCTCGAAAGCTCTGGTGTGCGCTACTGGCTGG |
| Comp-asymmetrical 36:90 | GTCTGGCAGCTCCCAATAAGGATCCCCCTTCCAGCCAGTAGCGCACACCAGAGCTTTCGAGAATGTTGATCACGTATTTAGTGGTCTCTCTGAGCGCACGTAAGTAGCAAGGTGGAGTCCAGCGCT |
| Rev Comp-asymmetrical 36:90 | AGCGCTGGACTCCACCTTGCTACTTACGTGCGCTCAGAGAGACCACTAAATACGTGATCAACATTCTCGAAAGCTCTGGTGTGCGCTACTGGCTGGAAGGGGGATCCTTATTGGGAGCTGCCAGAC |
| ***fkrp* C355Y with Guide 3 – 6 changes** | |
| Comp-asymmetrical 90:36 | TTTCGAGAATGTTGATCACGTATTTAGTGGTCTCTCGCAGCGCGCGCAGGTAGCAAGGTGGAGTCCAGCGCTCAATGTAAAGATATTCCGGTGTGTCATCTCGAACTGTCCCGAAACATCGCGCAG |
| Rev Comp-asymmetrical 90:36 | CTGCGCGATGTTTCGGGACAGTTCGAGATGACACACCGGAATATCTTTACATTGAGCGCTGGACTCCACCTTGCTACCTGCGCGCGCTGCGAGAGACCACTAAATACGTGATCAACATTCTCGAAA |
| Comp-symmetrical 60:60 | CCAGCCAGTAGCGCACACCAGAGCTTTCGAGAATGTTGATCACGTATTTAGTGGTCTCTCGCAGCGCGCGCAGGTAGCAAGGTGGAGTCCAGCGCTCAATGTAAAGATATTCCGGTGTGT |
| Rev Comp-symmetrical 60:60 | ACACACCGGAATATCTTTACATTGAGCGCTGGACTCCACCTTGCTACCTGCGCGCGCTGCGAGAGACCACTAAATACGTGATCAACATTCTCGAAAGCTCTGGTGTGCGCTACTGGCTGG |
| Comp-symmetrical 36:36 | TTTCGAGAATGTTGATCACGTATTTAGTGGTCTCTCGCAGCGCGCGCAGGTAGCAAGGTGGAGTCCAGCGCT |
| Rev Comp-symmetrical 36:36 | AGCGCTGGACTCCACCTTGCTACCTGCGCGCGCTGCGAGAGACCACTAAATACGTGATCAACATTCTCGAAA |
| Comp-asymmetrical 36:90 | GTCTGGCAGCTCCCAATAAGGATCCCCCTTCCAGCCAGTAGCGCACACCAGAGCTTTCGAGAATGTTGATCACGTATTTAGTGGTCTCTCGCAGCGCGCGCAGGTAGCAAGGTGGAGTCCAGCGCT |
| Rev Comp-asymmetrical 36:90 | AGCGCTGGACTCCACCTTGCTACCTGCGCGCGCTGCGAGAGACCACTAAATACGTGATCAACATTCTCGAAAGCTCTGGTGTGCGCTACTGGCTGGAAGGGGGATCCTTATTGGGAGCTGCCAGAC |

| ***fkrp* C355Y with Guide 1 – 1 change** | |
|---|---|
| Comp-asymmetrical 90:36 | GTATTTAGTGGTCTCTCTGAGGGCACGTAAGTAACAAGGTGGAGTCCAGCGCTCAATGTAAAGATATTCCGGTGTGTCATCTCGAACTGTCCCGAAACATCGCGCAGTCTCTTTACTGCAGCCAAA |
| Rev Comp-asymmetrical 90:36 | TTTGGCTGCAGTAAAGAGACTGCGCGATGTTTCGGGACAGTTCGAGATGACACACCGGAATATCTTTACATTGAGCGCTGGACTCCACCTTGTTACTTACGTGCCCTCAGAGAGACCACTAAATAC |
| Comp-symmetrical 60:60 | AGAGCTTTCGAGAATGTTGATCACGTATTTAGTGGTCTCTCTGAGGGCACGTAAGTAACAAGGTGGAGTCCAGCGCTCAATGTAAAGATATTCCGGTGTGTCATCTCGAACTGTCCCGAA |
| Rev Comp-symmetrical 60:60 | TTCGGGACAGTTCGAGATGACACACCGGAATATCTTTACATTGAGCGCTGGACTCCACCTTGTTACTTACGTGCCCTCAGAGAGACCACTAAATACGTGATCAACATTCTCGAAAGCTCT |
| Comp-symmetrical 36:36 | GTATTTAGTGGTCTCTCTGAGGGCACGTAAGTAACAAGGTGGAGTCCAGCGCTCAATGTAAAGATATTCCGG |
| Rev Comp-symmetrical 36:36 | CCGGAATATCTTTACATTGAGCGCTGGACTCCACCTTGTTACTTACGTGCCCTCAGAGAGACCACTAAATAC |
| Comp-asymmetrical 36:90 | GGATCCCCCTTCCAGCCAGTAGCGCACACCAGAGCTTTCGAGAATGTTGATCACGTATTTAGTGGTCTCTCTGAGGGCACGTAAGTAACAAGGTGGAGTCCAGCGCTCAATGTAAAGATATTCCGG |
| Rev Comp-asymmetrical 36:90 | CCGGAATATCTTTACATTGAGCGCTGGACTCCACCTTGTTACTTACGTGCCCTCAGAGAGACCACTAAATACGTGATCAACATTCTCGAAAGCTCTGGTGTGCGCTACTGGCTGGAAGGGGGATCC |

| ***nf1a* R628* with Guide 1 – 1 change** | |
|---|---|
| Comp-symmetrical 36:36 | CAGACACACCTCCAGCTTGGTCTGAGCCTGTCAGCTGATCGGGATCAGACTGCCCTGTGTAGCACACTCctg |
| Rev Comp-symmetrical 36:36 | cagGAGTGTGCTACACAGGGCAGTCTGATCCCGATCAGCTGACAGGCTCAGACCAAGCTGGAGGTGTGTCTG |

| ***nf1a* M992del with Guide 1 – 0 changes** | |
|---|---|
| Comp-symmetrical 36:36 | aaagaatagtaacaccttacCTAACTAGATTGAGCATCGTCTCAATGCTGGCTTGACCAAGGTGCTCTGAGC |
| Rev Comp-symmetrical 36:36 | GCTCAGAGCACCTTGGTCAAGCCAGCATTGAGACGATGCTCAATCTAGTTAGgtaaggtgttactattcttt |

| ***tp53 R144H* with Guide 1 – 0 changes** | |
|---|---|
| Comp-symmetrical 36:36 | cTATCTCCATCCGGGGTTCGCTCATGATGGGGGCAGTGGCGGACCACTTCAGCCACATGCTCGGACTTCTTA |
| Rev Comp-symmetrical 36:36 | TAAGAAGTCCGAGCATGTGGCTGAAGTGGTCCGCCACTGCCCCCATCATGAGCGAACCCCGGATGGAGATAg |

| ***pkd2* L593W with Guide 1 – 1 change** | |
|---|---|
| Comp-symmetrical 36:36 | ACACAAAGGTGGTGAAGTAAATGGGTCCCCAGACGCTGTCGGCCTCCTCGATTTCTGAGAAGTCGAAATCAC |
| Rev Comp-symmetrical 36:36 | GTGATTTCGACTTCTCAGAAATCGAGGAGGCCGACAGCGTCTGGGGACCCATTTACTTCACCACCTTTGTGT |

| ***tp53* K88R with Guide 1 – 1 change** | |
|---|---|
| Comp-asymmetrical 90:36 | actcacAGTGCAAGTTACAGATCTCGCTGTGCCAGACTGCGGGAACCTGAGCCTAAATCCATGATCGCCGGGATAGTCGCTTGTCTCCGGAACAGTGGATGTTGGTGG GAGAGTGGATGGCTGAGG |
| Rev Comp-asymmetrical 90:36 | CCTCAGCCATCCACTCTCCCACCAACATCCACTGTTCCGGAGACAAGCGACTATCCCGGCGATCATGGATTTAGGCTCAGGTTCCCGCAGTCTGGCACAGCGAGATC TGTAACTTGCACTgtgagt |
| Comp-asymmetrical 36:90 | tgaaatgtctctttccattaaatttgtgaaagtatgtgtgtgtatgcgcttttgactcacAGTGCAAGTTACAGATCTCGCTGTGCCAGACTGCGGGAACCTGAGCCTAAATCCATGATCGCCGGG |
| Rev Comp-asymmetrical 36:90 | CCCGGCGATCATGGATTTAGGCTCAGGTTCCCGCAGTCTGGCACAGCGAGATCTGTAACTTGCACTgtgagtcaaaagcgcatacacacacatactttcacaaatttaatggaaagagacatttc a |

### Supplemental Table 2

| | HRM forward | HRM reverse | NGS forward | NGS reverse | Sequencing forward | Sequencing reverse |
|---|---|---|---|---|---|---|
| *nf1a* R1191Q | CTCTTCGACTCTCGCCATCT | CTTCACGTCACCTTGAAGCA | CTCTTCGACTCTCGCCATCT | GAATGACTGTCACCGCTTCA | GTCATCGGAGTGCTGTTGTG | CTGCTGCCACTCTGGTGTAG |
| *nf1a* G763R | CCATTCATGGTCATCTATTTTCTG | AGGACTGTAGGTGCCCAGAC | CTGTTCGTCCTCAGGAGTGG | GGAGCTCATGGAAATCATGG | CATATGCAAAGTGGGAGCAA | AGTTCATGTCAAGCCCTGCT |
| *vma21* A58V | TGCAGTGTTTTTGATTTGCTTT | ATAAACGAAGAGCGCCAGAA | GGTGCAGTGTTTTTGATTTGC | ATAAACGAAGAGCGCCAGAA | CAAAATACGGGAGCTGTTGA | ATGCTTCCAGTGGGACATTC |
| *spop* D144N | CAAGGGAAAGACTGGGGTTT | TTAAACCGGTGATGGAAAGC | TTTTATGCAGAGAGCCAACG | TTAAACCGGTGATGGAAAGC | CAAGGGAAAGACTGGGGTTT | TAATGGCTGCTCAAAATTGC |
| *sgo1* K19E | CAGAAGAAGAGCTTCCAGCAG | AGACCGAGCGTCTCTCCAC | GACATACTGCGGGTGTTTGA | GTCTCTCCACCGACCTGCT | GTGAGTGAAAACAGCGGAGG | TTCAGCATCAACCAGACCGA |
| *pex10* H282D | CTCGCACTTCACGCTGTATC | TTCACAGCATCACTTCACAGC | CGCCGTTTTTCTCTTCAGTC | TTCACAGCATCACTTCACAGC | CGCCGTTTTTCTCTTCAGTC | TTCCATCACGATCACCTCAA |
| *fkrp* C355Y | ATGTTTCGGGACAGTTCGAG | AGTAGCGCACACCAGAGCTT | ATGTTTCGGGACAGTTCGAG | AATAAGGATCCCCCTTCCAG | TCATTCTCAGCCAGTCATCG | GAAATCGCCTTCAACAGCTC |
| *nf1a* R628* | agctcatttcctccacacag | GTACAGACACACCTCCAGCT | tgcccgtccagataaaacaag | CTGAAACAGGACATGGCCAC | ataacaattgcccgtccaga | actgacCTGTGGCCATCATA |
| *nf1a* M907del | AGGAAGCTCAGAGCACCTTG | tgaccatcagtctgtgtgacaa | ACTTGCTGGACAGTCACACT | tttatcaaaacaccaaccatcaa | cgtggtgccaactcagtttt | GCTCCATCATCACCTCCACT |
| *tp53* R144H | AAAACTTGCCCCGTTCAAAT | acTATCTCCATCCGGGGTTC | TCAAATGGTGGTGGACGTTG | gtgcaggcctagaatgatgc | tgctaaactataacaactgggtgaa | gtgcaggcctagaatgatgc |
| *pkd2* L593W | tttgctcacagTTTCACGCA | gtttacCAGCAGAATCATGAAGA | tttgctcacagTTTCACGCA | tgttttgtcattcatttcatgtaggg | tttgctcacagTTTCACGCA | CTGAGTTTTCCTGCTGCCTG |
| *tp53* K88R | tgtgtgtgtatgcgcttttg | TTCCGGAGACAAGCGACTAT | GTGCTTGAAGAACAGCCTCA | tgtgtgtgtatgcgcttttga | GAAGAACAGCCTCAGCCATC | accatggcacatatgcaaac |

| | ICE forward | ICE reverse | Allele Specific PCR forward | Allele Specific PCR reverse |
|---|---|---|---|---|
| *nf1a* R1191Q | CTCTTCGACTCTCGCCATCT | CTGCTGCCACTCTGGTGTAG | CAGACGCTCTTTCAAGGAAA | CTGCTGCCACTCTGGTGTAG |
| *nf1a* G763R | GCAAAGTGGGAGCAAGCTAC | AGCGATTCAGGGATGTTTCG | GGTTTCTTGTGCGCCTTAA | AGTTCATGTCAAGCCCTGCT |
| *vma21* A58V | CAAAATACGGGAGCTGTTGA | ATGCTTCCAGTGGGACATTC | TTTCTCTTGATAGTCGCCCTGG | GCCTCCACCATACCATGAGA |
| *spop* D144N | CAAGGGAAAGACTGGGGTTT | TAATGGCTGCTCAAAATTGC | TCTAAACGAAGCAAACGGGCT | TAATGGCTGCTCAAAATTGC |
| *sgo1* K19E | AGTGAAAACAGCGGAGGAGA | TGCAAGAGTAACAAGCTGTCA | GAGAAGATGGAAGAGAAGCGG | GCACATTTGGACTGTCTGCT |
| *pex10* H282D | CGCCGTTTTTCTCTTCAGTC | TTCCATCACGATCACCTCAA | CGGCGACCTGTTCTGCTG | TTCCATCACGATCACCTCAA |
| *fkrp* C355Y | ATGTTTCGGGACAGTTCGAG | GAAATCGCCTTCAACAGCTC | TTGCTACTTACGTGCCCTCA | GAAATCGCCTTCAACAGCTC |
